# Self-Supervised AI Discovery of Histomorphological Phenotypes from Routine Mesothelioma Biopsies

**DOI:** 10.64898/2026.08.09.743741

**Authors:** Farzaneh Seyedshahi, Francesca Damiola, Ruth Sequeiros, Fabien Forest, Arnaud Scherpereel, Ke Yuan, Sylvie Lantuejoul, John Le Quesne

**Affiliations:** School of Cancer Sciences, University of Glasgow, Glasgow, UK; Cancer Research UK Scotland Institute, Glasgow, UK; National Reference Center MESOPATH-NETMESO and French MESOBANK, Biopathology Department, Centre Léon Bérard, Lyon, France; Department of Biopathology, Saint-Étienne University Hospital (Hôpital Nord), Saint-Étienne, France; Saint-Étienne University, Saint-Étienne, France; CHU de Lille, Department of Pneumology and Thoracic Oncology; NETMESO expert network, Lille, France; University of Lille; INSERM U1189 - ONCO-THAI - Image Assisted Laser Therapy for Oncology, ONCOLille, Lille, France; Grenoble Alpes University, Grenoble, France; Queen Elizabeth University Hospital, NHS Greater Glasgow and Clyde, Glasgow, Scotland, UK

## Abstract

Accurate subtype diagnosis is essential for guiding therapy and predicting patient outcome in malignant mesothelioma. Most computational pathology models are trained on large tissue images from resection specimens, which maximises information for training but limits model relevance in real-world diagnostic settings where small biopsies are the most usual tissue source. In this work, we assembled a large multicentre cohort of mesothelioma biopsy slides. We used a self-supervised learning model to evaluate the associations of biopsy-driven morphology patterns with histological subtype, molecular markers, and survival. The discovered histomorphology patterns captured a continuum of tissue phenotypes spanning epithelioid and sarcomatoid malignancy and non-tumour morphologies. Furthermore, patient-level HPC representations performed excellently in distinguishing epithelioid from non-epithelioid mesothelioma (AUC = 0.94) and demonstrated prediction of immunohistochemistry (IHC) markers. Additionally, HPC-derived features alone achieved performance comparable to established clinical and molecular variables (C-index = 0.65), while integration of HPCs with clinical and marker information improved performance to a C-index of 0.69. Several HPCs were significantly associated with either favourable or adverse prognosis and reflected both clinically known and biologically plausible patterns. In conclusion, self-supervised learning can discover interpretable histomorphological phenotypes directly from routine mesothelioma small biopsies without further training. These AI-derived phenotypes capture clinically and biologically relevant information, linking tissue architecture to molecular characteristics, histological subtypes, and patient outcomes. The proposed framework provides a thorough evaluation of real-world biopsy data using a pre-trained model, without the need for computationally intensive retraining, and addresses the question of whether SSL-based AI can be deployed out of the box in clinical settings.

## 2 Main

Mesothelioma is a rare but aggressive cancer, almost universally caused by asbestos exposure, and characterised by profound histological heterogeneity and poor prognosis ^1, 2^. Accurate histo-subtype classification, especially into epithelioid and non-epithelioid categories is essential for accurate prognosis and to guide therapeutic decisions. However, current expert histopathological diagnosis remains subjective and can suffer from high inter-observer variability, particularly in the diagnosis of difficult “edge case” appearances ^3, 4, 5^. Immunohistochemistry (IHC) is a vital tool for mesothelioma diagnosis ^6, 7^, and markers including Wilms’ Tumor 1 (WT1), Cytokeratin 5/6 (CK5/6), Ber-EP4, BRCA1 Associated Protein-1 (BAP1), Calretinin, Thyroid transcription factor-1 (TTF-1), and Cytokeratin AE1/AE3 are among the effective IHC markers for mesothelioma identification or exclusion ^8, 9, 10, 11, 12, 13^.

Recent advances in computational pathology have shown promise in addressing histopathological challenges in mesothelioma. For instance, weakly-supervised deep-learning models such as MesoNet ^14^ have been used to predict overall survival directly from whole-slide images, outperforming standard pathology practice while also uncovering region-level prognostic features rooted in inflammation and stromal architecture. However, the method relies heavily on human gold-standard subtype labels to train the encoder model to recognize morphological patterns, limiting model performance to human expertise, and requiring substantial time investment prior to training. More recently, MesoGraph ^15^, a graph neural network (GNN) framework, enabled cell-level scoring of sarcomatoid and epithelioid phenotypes using a small subset of tissue microarrays (TMAs). It produces a continuous MesoScore that captures both histological subtype composition and patient survival risk. However, GNN-based approaches are often not scalable or computationally practical for larger tissue samples, such as resections and biopsies, and tend to break down in real-world clinical settings ^16^. Additionally, training procedures such as MesoNet rely on core-level subtype annotations within a weakly supervised framework. As a result, the cellular and, subsequently, core-level representations learned by these models remain strongly anchored to human-provided labels rather than directly capturing underlying tissue morphology. This dependence is particularly limiting in mesothelioma, a disease characterised by substantial inter-pathologist disagreement and ambiguous histomorphological patterns. Consequently, label noise may propagate through the model, reducing robustness and potentially obscuring clinically meaningful morphological signals.

In response to the limitations of weakly supervised learning, self-supervised learning (SSL) approaches have recently gained substantial traction in computational pathology. Large-scale foundation models such as UNI ^17^, Prov-GigaPath ^18^, CONCH^19^, Virchow^20^, MUSK^21^, and others have demonstrated the ability to learn robust, transferable morphological representations directly from unlabelled histopathology images, substantially reducing reliance on potentially noisy human annotations. By decoupling representation learning from diagnostic labels, these methods aim to mitigate human-induced bias and improve generalisability across centres and disease contexts. In mesothelioma, a recent study used Histomorphology Phenotype Learning (HPL) ^22^, a self-supervised approach trained on surgically resected tissue to generate a histomorphological atlas capturing recurrent phenotypes across thousands of wole-slide images ^23^. This atlas demonstrated clinically meaningful performance for both prognostic stratification and subtype classification. In this work, we showed that training a self-supervised deep learning feature encoder on a large-scale dataset of nearly 2.5 million tiles enables the development of a mesothelioma-specific foundation model that more faithfully captures the disease’s underlying morphology.

I real-world practice, mesothelioma diagnosis and prognostic assessment rely mainly on small tissue biopsies; however, most computational pathology models, including HPL-Meso ^23^, are trained on large tissue samples from surgical resections, thereby maximising training data. However, these large resection specimens contain extensive highly morphologically diverse amounts of tumour, stroma, necrosis, inflammation, and background tissue image areas, whereas by comparison, biopsies are much smaller, more uniform in composition, and are prone to different processing artefacts compared to resections (eg. more crush artefact, more over- vs under-fixation), and face sampling limitations. These differences create a significant mismatch between the WSI data used for model development and the biopsy material used in most diagnostic workflows, with direct implications for model reliability and clinical applicability.

To address this gap, we assembled a large multicenter French biopsy cohort comprising approximately 1,000 patients and samples from more than 100 institutions along with corresponding IHC data and diagnoses from NETMESO Mesopath panel pathologist reviews. In contrast to the routine UK hematoxylin and eosin (H&E) staining protocol, the French material, retrieved from the Mesobank.fr collection, was stained with HPS (hematoxylin–phloxine–saffron) and HES (hematoxylin–eosin–saffron) as per the national standard, giving an additional yellow/colour to dense collagenous tissue. This French national dataset represents an extraordinary learning resource, as most if not all other cohorts are smaller and fragmented, and lack such comprehensive IHC data and central review.

We then evaluated the exisiting HPL-Meso model which had previously been trained on UK surgical resection WSIs stained with hematoxylin and eosin (H&E) ^23^. The UK-trained image encoder was applied to the HES/HPS-stained French biopsy dataset to extract mesothelioma-specific features previously discovered in resections, thereby applying learnings from one dataset to another markedly divergent in both sample type and staining protocol. The setting introduces two major domain shifts: (a) a change in sample type and extent (resection to biopsy) and (b) staining differences due to the presence of saffron-based collagen highlighting and phloxine contrast. Additional variations in institution, country, and scanning protocols further increase the real-world clinical relevance of this work.

The key open question is whether self-supervised encoders trained on large resections can be reliably transferred to biopsy material while retaining diagnostic fidelity. To isolate this domain shift within a clinically realistic mesothelioma setting, we deliberately restrict the experimental framework to this single cancer and a fixed encoder architecture, avoiding confounding effects introduced by cross-cancer generalisation or model heterogeneity. Within this controlled setting, we evaluate the extent to which the pretrained encoder maintains diagnostically and prognostically relevant signals when applied to real-world clinical biopsies, without further training. We then assess the utility of biopsy-derived histomorphological clusters for patient-level survival prediction and for accurate classification of epithelioid versus non-epithelioid disease subcategories.

## 3 Results

### 3.1 Evaluation across a Multi-Center French Biopsy Cohort

For this study, we collected a large multicentre cohort of HES- and HPS-stained biopsy slides received from pathology laboratories across several regions in France. All slides were reviewed by at least three subspecialty mesothlioma expert pathologists of the French NETMESO MESOPATH network, and digitised at the National Reference Centre in Lyon using Leica AT2 or Leica GT scanners at 20X magnification. The final dataset comprised 1,047 patients and 1,062 biopsy images. Among the patients, 77% (*n* = 806) were male and 23% (*n* = 241) were female. Patient ages ranged from 22 to 100 years, with a mean age of 76.5 ± 8.7 years (Table 1). Treatment information was available for 199 patients, of whom 108 received chemotherapy, 53 radiotherapy, 21 surgery, and 17 immunotherapy. Both age and mesothelioma subtype were significantly (*p <* 0.05) associated with survival (Supplementary Figure S1). The cohort included 61.6% epithelioid (*n* = 645) and 38.4% non-epithelioid (*n* = 402) cases. Figure 1a illustrates the geographical distribution of referring institutes and highlights the top 10 contributing cities, led by Lyon. The map partly reflects differences in the number and capacity of pathology laboratories between regions.

**Table 1.**
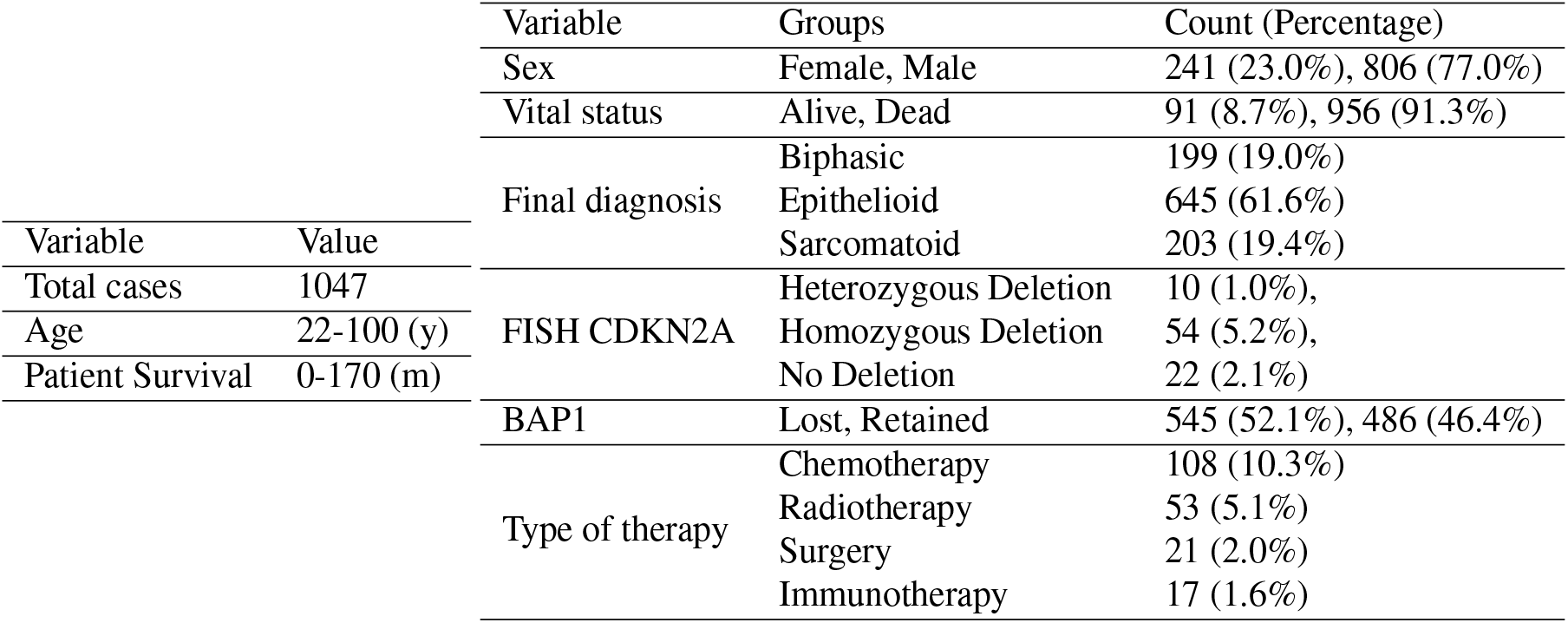
Dataset Characteristics.

| Variable | Value | Variable | Groups | Count (Percentage) |
| --- | --- | --- | --- | --- |
| Total cases | 1047 | Sex | Female, Male | 241 (23.0%), 806 (77.0%) |
| Age | 22-100 (y) | Vital status | Alive, Dead | 91 (8.7%), 956 (91.3%) |
| Patient Survival | 0-170 (m) | Final diagnosis | Biphasic | 199 (19.0%) |
|  |  |  | Epithelioid | 645 (61.6%) |
|  |  |  | Sarcomatoid | 203 (19.4%) |
|  |  | FISH CDKN2A | Heterozygous Deletion | 10 (1.0%), |
|  |  |  | Homozygous Deletion | 54 (5.2%), |
|  |  |  | No Deletion | 22 (2.1%) |
|  |  | BAP1 | Lost, Retained | 545 (52.1%), 486 (46.4%) |
|  |  | Type of therapy | Chemotherapy | 108 (10.3%) |
|  |  |  | Radiotherapy | 53 (5.1%) |
|  |  |  | Surgery | 21 (2.0%) |
|  |  |  | Immunotherapy | 17 (1.6%) |

**Figure 1.**
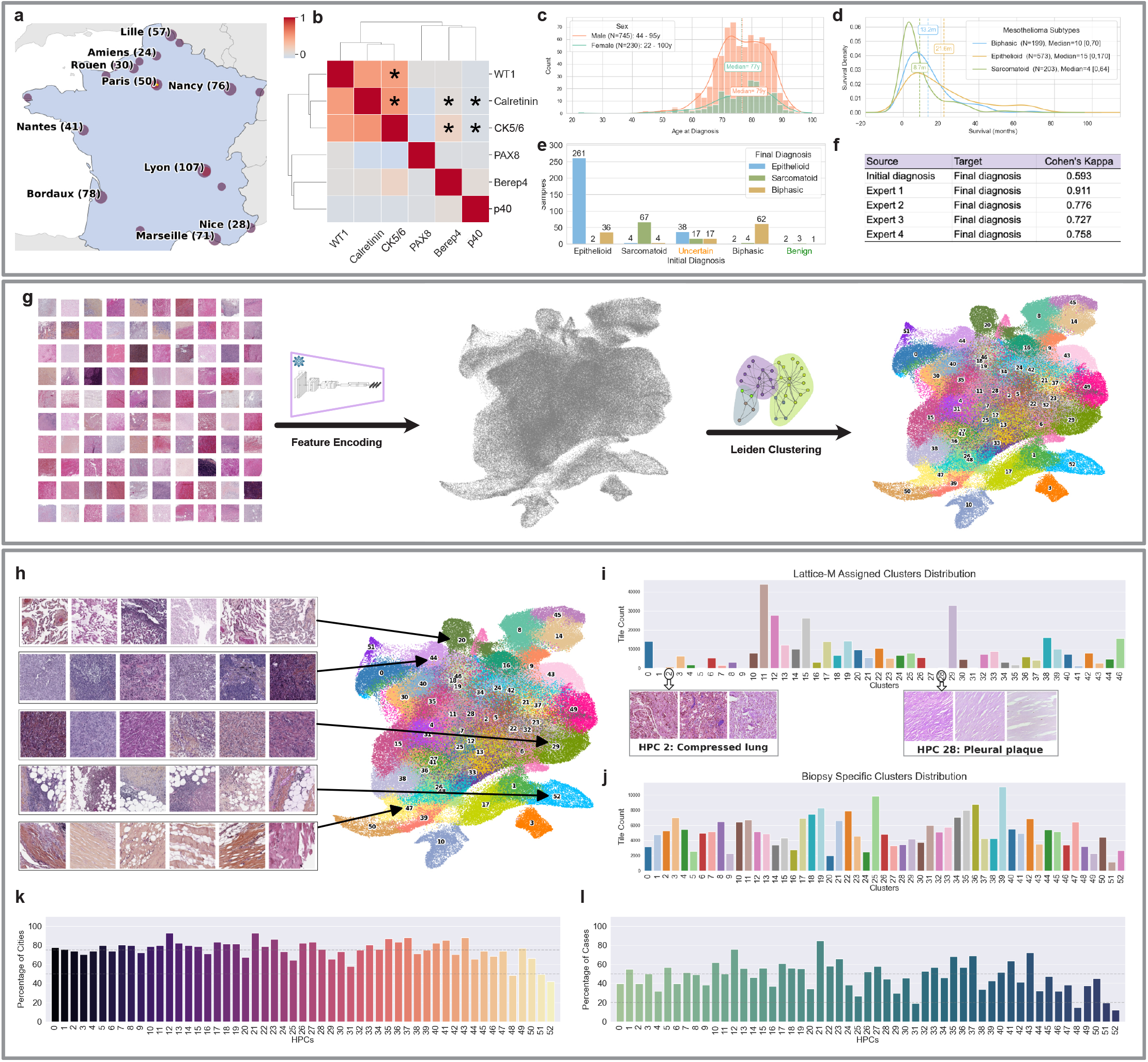
Overview of the multicentre mesothelioma biopsy cohort, immunohistochemical marker relationships, diagnostic variability, and histomorphological phenotype clusters (HPCs). **(a)** Geographical distribution of referring pathology centres and patient case numbers across French cities. **(b)** Pairwise correlation matrix of immunohistochemistry (IHC) markers, with significant correlations indicated by asterisks (p<0.01). **(c–d)** Distribution of patient age at diagnosis and survival across mesothelioma subtypes. **(e–f)** Diagnostic variability analysis showing changes between initial and final diagnoses and agreement scores between pathologists and consensus diagnosis. **(g–h)** Workflow for generating biopsy-derived HPCs and UMAP visualisation of tile embeddings coloured by cluster assignment. Representative clusters illustrate distinct morphological patterns, including alveolar tissue, inflamed and highly cellular tumorous tissue, and adipose tissue. **(i–j)** Comparison between resection-derived and biopsy-specific HPC assignments, highlighting the limited representation of certain resection-associated morphologies in biopsy samples. **(k–l)** Geographical and patient-level distribution of biopsy-derived HPCs, demonstrating broad representation of clusters across centres and cases.

In Figure 1b, we computed pairwise correlations between tumour cell immunohistochemistry markers available in more than 100 samples. Significant correlations (*p <* 0.01) are indicated with an asterisk. The analysis focused on CK5/6, calretinin, and WT1, which were used as positive markers for mesothelioma, and BerEP4, TTF-1, p40, and PAX8, which were used as negative markers. CK5/6, calretinin, and WT1 showed strong positive correlations with one another. Although p40 (and PAX8) expression was largely absent or focal in this mesothelioma cohort, it was included in the figure to illustrate its absence in relation to the other markers. A more complete summary of reported marker distributions is provided in Supplementary Figure S2.

The median age at diagnosis for male patients was 79 years (minimum 44 years), compared with 77 years for female patients (minimum 22 years) (Figure 1c). Survival distributions differed substantially across mesothelioma subtypes, with mean survival times of 8.7, 13.2, and 21.6 months for sarcomatoid, biphasic, and epithelioid cases, respectively. Sarcomatoid cases exhibited the poorest prognosis, with a median survival of only 4 months, compared with 10 months for biphasic and 15 months for epithelioid cases (Figure 1d).

To highlight the diagnostic challenges associated with mesothelioma, Figure 1e shows that 36 patients initially diagnosed with epithelioid disease were later reclassified as biphasic following central review by the French national board of experts. Among 72 cases with uncertain initial diagnoses, the final diagnoses were given as epithelioid (*n* = 38), sarcomatoid (*n* = 17), and biphasic (*n* = 17). In addition, three cases initially considered benign were ultimately classified as sarcomatoid malignancy, further emphasising the practical value of expert board review. Indeed, the Cohen’s kappa score between initial and final diagnoses was 0.59, indicating only moderate agreement. Agreement scores between individual experts and the final consensus diagnosis ranged from 0.73 to 0.91 (Figure 1f).

### 3.2 Discovery of Patient-Agnostic Histomorphological Phenotype Clusters (HPCs)

To characterise morphological patterns across the cohort, we used the resection specimen-trained feature encoder from ^23^ to generate tile-level feature embeddings and construct histomorphological phenotype clusters (HPCs) (Figure 1g). Leiden clustering identified 53 distinct HPCs. The UMAP projection of tile embeddings is shown in Figure 1h, with tile embeddings coloured according to cluster assignment. Although the encoder was trained exclusively on H&E images, it remained robust to staining variation and generated discriminative feature representations despite the presence of saffron staining. Clusters demonstrated clear morphological distinctions. For example: cluster 20 exhibited pulmonary alveolar architecture; clusters 44 and 29 displayed solid/trabecular pattern malignancy, with and without dense chronic inflammation, respectively; cluster 52 was characterised by inflamed and fibrotic adipose tissue from the chest wall; and cluster 47 corresponded to benign fibroadipose and artefact tissues.

As shown in Figure 1i-j, assigning biopsy tile embeddings to previously defined resection-derived HPCs (from the HPL-Meso study ^23^) constrained the representation space to the morphological vocabulary of surgical resection samples. As might be expected, several essentially resection-specific HPCs, such as HPC 28 (pleural plaque) and HPC 2 (compressed lung), were absent in biopsy specimens, and others were over-represented. Therefore we delineated a new set of biopsy-specific HPCs from the French cohort’s latent representations. These HPCs exhibited a substantially more balanced distribution of tiles, and were used for all downstream analyses. The distribution of HPC contributions across cities is shown in Figure 1k and demonstrates a good geographical balance and diversity, and no HPC belongs to a single institute. Every HPC contained biopsy tiles originating from at least 42% of represented cities, while 50 of the 53 clusters received contributions from more than 50% of cities. Among the 107 represented cities, cluster 52 showed the lowest geographical coverage, with contributions from 45 cities, whereas cluster 12 showed the broadest coverage, with tiles originating from 99 cities. Figure 1l further illustrates the distribution of HPCs across individual patients. Of the 53 identified HPCs, 49 were represented in at least 20% of cases, while 23 HPCs were observed in more than 50% of cases, suggesting the presence of recurrent and common mesothelioma morphological patterns across the cohort.

### 3.3 Pathologist Validation and Alignment with Immunohistochemistry Profiles

After defining the HPC set, we aimed to assign morphological meaning to each cluster and better characterise the histomorphological backbone of the cohort. Three subspecialty expert pathologists independently reviewed 100 representative random tiles from each HPC and annotated a range of morphological features, including necrosis, inflammation, architectural pattern, nuclear atypia, and stromal cellularity. Figure 2a summarises the consensus subtype labels assigned to each HPC. HPCs with complete agreement between all three experts (3/3 same annotations) are marked with an asterisk (*), whereas HPCs with partial agreement (2/3) display the majority-vote subtype label. In cases of no agreement, observed for HPCs 30 and 45 (2/53, 3.8%), no consensus subtype label was assigned. The average pairwise inter-observer agreement across all HPCs was moderate (*κ* = 0.584, *n* = 53). Full agreement among all three experts was achieved for 32/53 (60.4%) HPCs, while partial agreement was observed for 19/53 (35.8%).

**Figure 2.**
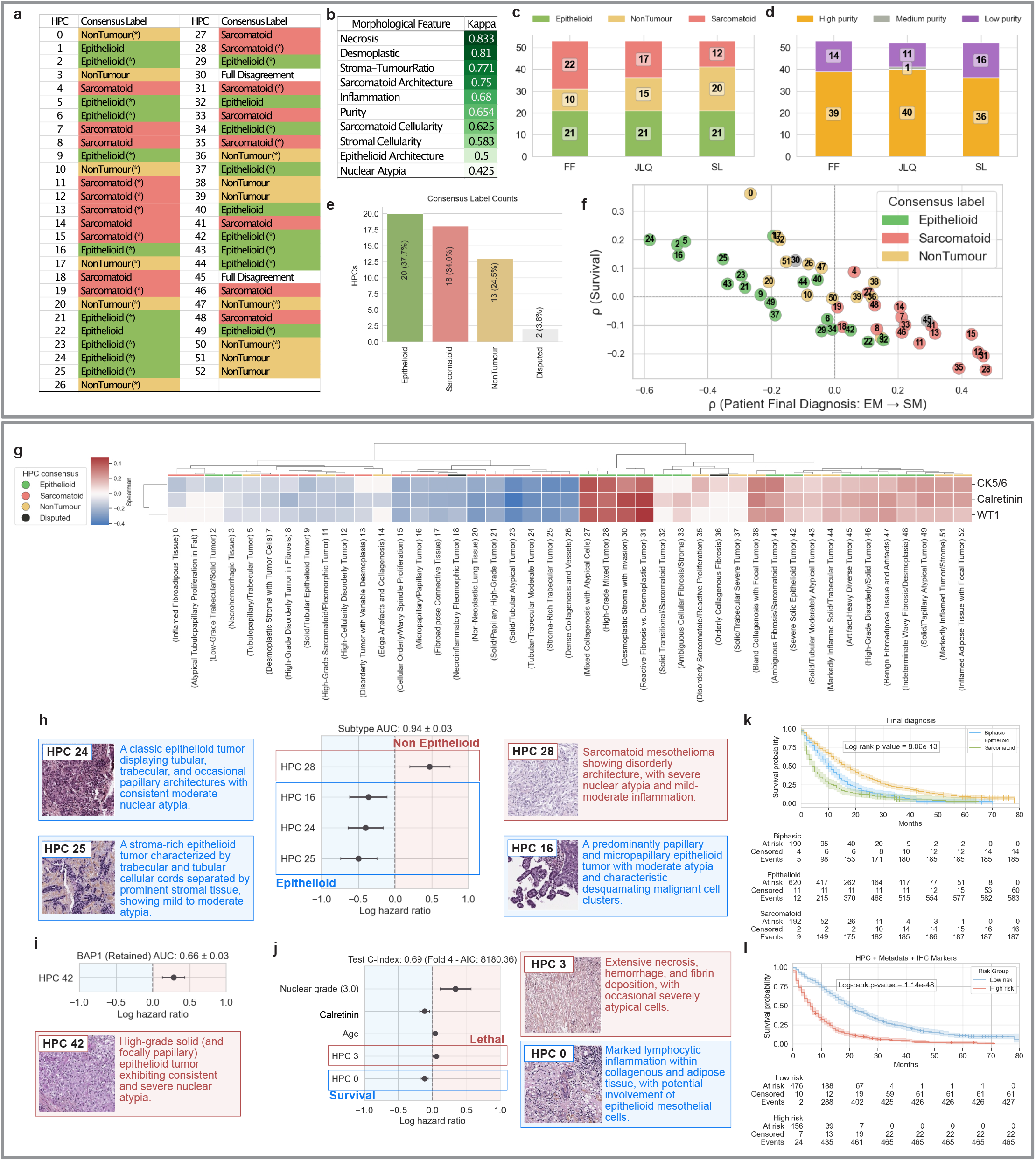
HPC Annotations and downstream tasks (a-f) HPC annotations’ analysis (g)Significant IHC marker correlations with HPCs. (h-j) Significant HPCs associated with downstream tasks. (K,I) Kaplan-Meier survival curves stratified by histological subtype and by integrated risk groups derived from clinical, molecular, and HPC features.

Among the annotated histomorphological features, necrosis showed the highest inter-observer agreement, with a Cohen’s kappa score of 0.833, whereas nuclear atypia demonstrated the lowest agreement (*κ* = 0.425). Inflammation scoring also showed substantial consistency between experts, with a kappa score of 0.68 (Figure 2b). Figures 2c and d present the distribution of epithelioid, sarcomatoid, and non-tumour categories, together with the estimated purity of each HPC (ie the degree to which the entire tile set shares common pathological features) according to the individual expert annotations. Overall, 20/53 (37.7%) HPCs were classified as epithelioid, 18/53 (34.0%) as sarcomatoid, and 13/53 (24.5%) as non-tumour, while 2/53 (3.8%) remained unclassified because of disagreement between reviewers (Figure 2e).

Using the occurrence frequency of each HPC across patient samples, we generated case-level embedding vectors following centred log-ratio (CLR) transformation to normalise and centralise the compositional data. Correlating individual HPC features with both survival duration and diagnostic subtype revealed distinct morphological trends across the cohort. Figure 2f illustrates these associations in a scatter plot, where each HPC is coloured according to its consensus subtype label. A gradual transition was observed from epithelioid-associated HPCs, which were positively correlated with survival and enriched in epithelioid cases, through non-tumour HPCs with weaker associations, to sarcomatoid-associated HPCs showing strong negative correlations with survival duration and positive association with sarcomatoid phenotype.

Figure 2g shows the significant correlations between CLR-transformed HPC embeddings and immunohistochemical markers. Statistical significance was assessed using FDR-corrected p-values (*p <* 0.05), and each HPC was labelled according to the expert consensus annotations. The classical diagnostic markers of mesothelioma, Calretinin, CK5/6 and WT1, identify a set of HPCs dominated by malignant epithelioid morphology, while sarcomatoid clusters are notably related to the loss of these markers.

### 3.4 HPCs Predict Core Mesothelioma Subtypes, BAP1 Loss, and Tumor Grade

Using the CLR-transformed HPC frequency vectors as case-level representations, we evaluated the ability of the learned histomorphological features to predict multiple downstream clinical and molecular variables, including histological subtype (epithelioid vs non-epithelioid), initial diagnosis, sarcomatoid versus biphasic subtype, BAP1 loss status, nuclear grade, and immunohistochemistry marker expression such as CK5/6 and Calretinin. Performance metrics, including Area under the curve (AUC), F1-score, sensitivity, and precision, are summarised in Table 2.

**Table 2.**
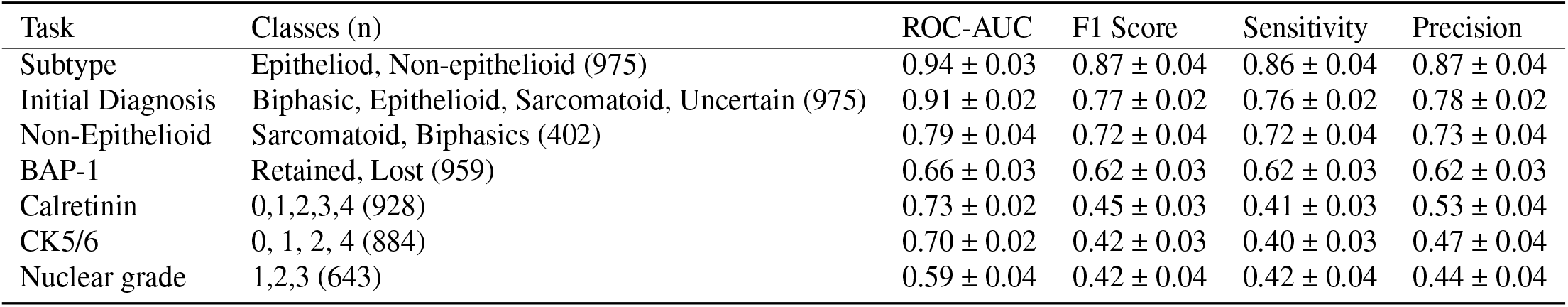
Performance of downstream classification tasks, including histological subtype, BAP1 loss, nuclear grading, and marker expression prediction using CLR-transformed HPC embeddings.

The framework achieved a cross-validated average AUC of 0.94 for distinguishing epithelioid from non-epithelioid cases (*n* = 975), demonstrating that the unsupervised HPC representations captured highly discriminative subtype-associated morphology. HPCs 16 (papillary architecture), 24 (tubular architecture), and 25 (trabecular architecture), all characterised by moderate nuclear atypia and minimal inflammation, were among the most significant predictors of epithelioid disease. In contrast, HPC 28, characterised by disorderly sarcomatoid architecture and severe nuclear atypia, was the strongest predictor of non-epithelioid cases (Figure 2h).

These findings indicate that the HPC representation space captures biologically meaningful architectural transitions associated with mesothelioma subtype progression.

The model also demonstrated predictive capability for BAP1 loss, achieving an average AUC of 0.66 with a sensitivity of 0.62 (*n* = 959). HPC 42, characterised by epithelioid solid architecture with severe nuclear atypia and mild-to-moderate inflammation, was significantly associated with BAP1 loss in case-level predictions. This observation is consistent with previous studies reporting higher frequencies of BAP1 loss in epithelioid mesothelioma compared with other histological subtypes ^24, 25^. Overall, these results suggest that the self-supervised HPC representations encode not only morphological subtype information, but also clinically relevant molecular and phenotypic characteristics directly from routine biopsy images.

### 3.5 High Grade Tumour Histological Phenotypes Drive Patient Risk

We next evaluated the prognostic value of the learned HPC information using survival analysis. Clinical variables, including age, sex, and histological subtype diagnosis (epithelioid, sarcomatoid, and biphasic), achieved an average test-set concordance index (C-index) of 0.65. In comparison, immunohistochemistry markers, including Calretinin, CK5/6, and WT1 (positive markers), demonstrated a predictive value with a 5 fold cross validation average C-index of 0.61 for the test set.

Incorporating histomorphological information derived from HPC statistics (such as Pielou’s evenness index and Shannon diversity) alongside their frequency of occurrence in the patient samples increased the test-set C-index to 0.65 (Table 3). Filtering HPCs using pathologist annotations by restricting the analysis to tumour-containing HPCs (*n* = 38) or high-purity HPCs (*n* = 40) did not improve the C-index. However, combining the HPC-derived frequencies and statistical metrics with both IHC data from positive markers and clinical information significantly increased performance to a cross-validated average C-index of 0.68.

**Table 3.**
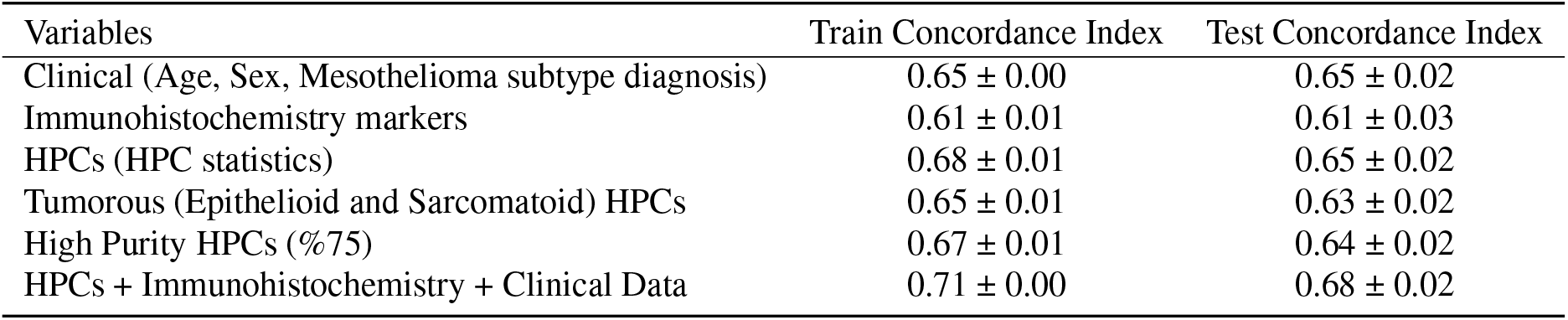
Survival prediction performance using clinical variables, immunohistochemistry markers, and HPC-derived features individually and in combination. Reported metrics are averaged across 5-fold cross-validation and are presented as mean ± standard deviation.

| Variables | Train Concordance Index | Test Concordance Index |
| --- | --- | --- |
| Clinical (Age, Sex, Mesothelioma subtype diagnosis) | 0.65 ± 0.00 | 0.65 ± 0.02 |
| Immunohistochemistry markers | 0.61 ± 0.01 | 0.61 ± 0.03 |
| HPCs (HPC statistics) | 0.68 ± 0.01 | 0.65 ± 0.02 |
| Tumorous (Epithelioid and Sarcomatoid) HPCs | 0.65 ± 0.01 | 0.63 ± 0.02 |
| High Purity HPCs (%75) | 0.67 ± 0.01 | 0.64 ± 0.02 |
| HPCs + Immunohistochemistry + Clinical Data | 0.71 ± 0.00 | 0.68 ± 0.02 |

These findings show that the self-supervised HPC representations capture prognostically relevant morphological patterns comparable in predictive strength to established clinical and immunohistochemical variables, which is remarkable considering the size of the biopsies and the complex polymorbidity faced by these patients. Moreover, the improved performance achieved through feature integration indicates that HPC-derived morphology provides complementary information rather than merely encoding disease subtype.

Figure 2j summarises the variables significantly associated with patient outcome. HPC 0, a cluster of mixed appearance characterised by marked inflammation was associated with relatively good outcome, perhaps indicating an effective antitumour immune response. In contrast, HPC3 predicts poor outcome; this cluster contains homogenous paucicellular tiles, many of which are wholly necrotic, which has previously been reported as a predictor of poor outcome ^26^. Additionally, older patient age and higher nuclear grade were significantly associated with an increased risk of death. Conversely, Calretinin expression was associated with improved survival. These findings further support the biological relevance of HPC-derived morphological representations and their alignment with established prognostic markers in mesothelioma.

Figure 2k demonstrates significant survival stratification across the three histological subtypes (*p* = 8.6 × 10^−13^). Furthermore, integrating clinical metadata, immunohistochemistry markers, and HPC statistics enabled stratification of patients into high-risk and low-risk groups with substantially stronger separation (*p* = 1.14 × 10^−48^; Figure 2l). This highlights the added prognostic value of quantitative histomorphological features beyond conventional subtype diagnosis alone and suggests that AI-derived tissue representations may provide a more nuanced characterisation of tumour behaviour and patient outcome.

## 4 Discussion

In this study, we evaluated the HPL-Meso encoder originally trained on H&E surgical resections from the University Hospitals of Leicester, UK (LATTICe-M) on a heterogeneous cohort of HPS and HES biopsy slides collected across multiple histopathology centres in France. Our goal was to derive a new dictionary of biopsy-specific histomorphological phenotype clusters, and assess the utility of these features for downstream clinical tasks, including survival prediction and mesothelioma subtype classification, both of which have direct diagnostic and therapeutic impact in this disease.

The decision to reuse a mesothelioma-trained encoder, rather than a multi-cancer foundation model, was intentional. We aimed for a model specific enough to capture disease-relevant morphology while remaining generalisable across institutions and staining protocols. Training a bespoke model from scratch, particularly in clinical environments with limited computational resources, is rarely feasible, making pretrained encoders an essential component of practical digital pathology pipelines.

The geographical analysis demonstrates strong spatial mixing across clusters, with no cluster dominated by a single diagnostic location, indicating that the learned histomorphological clusters are not driven by centre-specific artefacts or practices. Despite substantial variation in patient counts across cities, partly reflecting differences in the number and capacity of pathology laboratories, most clusters receive contributions from a majority of cities, supporting their robustness in a real-world, multi-centre setting. Importantly, the recorded location corresponds to the patient’s diagnostic centre rather than the slide preparation or digitisation site, and diagnostic workflows may involve multiple scans per patient, including secondary reviews by regional expert pathologists. As a result, it is not possible to determine whether the exact slide used for model training was the primary diagnostic slide; however, most slides, particularly HES/HPS, were reviewed by the initial pathologists. Also, training slides were drawn from those reviewed by expert pathologists during diagnosis, and the observed geographical diversity across clusters suggests limited impact of this uncertainty on cluster composition.

The key focus of this evaluation is the presence of two concurrent domain shifts: the application of a resection-trained model to biopsy material, and the transition from H&E to HPS/HES staining; the latter introducing saffron to enhance collagen visualisation. Given that saffron-based stains are routinely used in France and French-speaking Canada but remain uncommon elsewhere, this setting provides a rigorous context in which to examine the model’s capacity to generalise across both tissue substrate and staining practice. Additionally, scanner and colour profiles differed from those used to generate the primary training dataset (surgery resection LATTICe-M cohort). Despite this, the self-supervised encoder demonstrated strong colour invariance and successfully produced discriminative feature embeddings that organised biopsy tiles into well-separated phenotypic clusters. These clusters captured recognisable mesothelioma morphologies and were sufficiently stable to support downstream inference.

The clusters enabled meaningful survival modelling and delivered particularly strong performance in subtype classification (Table 2). The superior performance in the subtyping task (in comparison to LATTICe-M) likely reflects the nature of biopsy tissue; biopsies are collected with the intention of being near-pure samples of the tumour, reducing morphological heterogeneity and making overall cellular architecture more diagnostic. Survival prediction was inherently more challenging than subtype classification, as biopsies sample only a minute fraction of the tumour, greatly constraining the biological snapshot of the tumour mass as a whole. This restricted context naturally reduces prognostic signal compared to resection-based analyses. Nevertheless, a concordance index in the range of 0.65 - 0.70 should not be considered modest; within the literature^14^, such performance is regarded as strong when derived exclusively from histological images, without incorporation of clinical variables or patient history. This study used a higher magnification (40x) than we did but obtained similar scores. This suggests that having access to finer cell details cannot explain the results, though magnification could still be a factor. Aggregating multiple biopsy samples per patient, when available, is a promising strategy to recover missing architectural context and improve prognostic robustness. Despite this limitation, the learned representations provide sufficient discriminative information to robustly stratify patients into high- and low-risk groups, as reflected by significant survival separation. Furthermore, the correspondence between clusters derived from the LATTICe-M resections and biopsy-specific clusters indicates that a substantial portion of the biological dictionary learned from resections remains relevant across tissue types. This supports two key observations: first, that resection-derived morphological patterns capture universal tumour biology applicable to biopsies with meaningful prognostic coverage; and second, that the discovery of biopsy-specific clusters enables finer-grained characterisation, as certain resection-derived clusters are absent in biopsies and vice versa (Figure1i,j), ultimately expanding the morphological spectrum.

In summary, our results show that models trained on resection specimens still achieve clinically meaningful accuracy when applied to biopsy material. This validation also strengthens the evidence for the robustness of the HPL pipeline. Furthermore, despite being acquired without any expert or diagnostic supervision, the HPC features showed associations with molecular characteristics and immunohistochemical marker expression. Importantly, HPC-derived features provided prognostic information comparable to established clinical and molecular variables and further improved survival prediction when integrated with them. The identified HPCs corresponded to interpretable architectural and cytological patterns recognised by expert pathologists, suggesting that the learned representation space reflects genuine disease morphology rather than dataset-specific visual signals.

One limitation of this study is the incomplete transparency around the provenance of the external biopsy cohorts. Although we validated performance across two independent institutions and confirmed consistent results across the major scanner models used, we did not have access to fine-grained, site-specific metadata such as detailed instrument maintenance logs, reagent batch information, or full IT infrastructure records. This limits our ability to fully rule out subtle site-specific confounders and reflects an ongoing challenge in multi-institutional translational research.

Another limitation is that although we demonstrate that histology-derived phenotype features alone can support both diagnostic and prognostic tasks, the current framework does not integrate multimodal data such as next-generation sequencing or other genomic markers to train the model. The analysis relied exclusively on histomorphological signals, and incorporating genomic features will be an important next step to quantify the added value of computational pathology features over established clinical and molecular diagnostics. This multimodal extension training is already planned as a follow-up to the present work.

From a translational perspective, this validation framework offers a practical blueprint for integrating self-supervised pathology models into clinical workflows. It shows that pretrained, disease-specific encoders can be effectively repurposed for diagnostic stratification and risk assessment directly from routine biopsy slides, providing a scalable path toward AI-assisted mesothelioma diagnosis and clinical decision-making. While domain-specific differences inevitably introduce some uncertainty, systematic external validation on real-world biopsy data remains essential before clinical deployment.

To promote transparency and enable full reproducibility, we commit to releasing the biopsy-specific HPCs, upon request. The code for compositional feature extraction, and the trained SSL encoder weights are on public repository ^27^.

## 5 Methodology

To address the proposed research question, we designed a controlled experimental framework centered on mesothelioma biopsy material. The Methodology section is structured to first describe the datasets used, including the resection cohorts employed for self-supervised pretraining and the independent biopsy cohort.

We then outline the overall model pipeline, detailing the transfer of the pretrained encoder to biopsy slides, the extraction and clustering of tile-level representations, and the construction of patient-level features. Finally, we describe the experimental protocols used to assess diagnostic fidelity and prognostic relevance, with all quantitative results reported in the Results section.

The feature encoder used in this work was trained with the Barlow Twins SSL framework on the Leicester Archival Thoracic Tumour Investigation Cohort–Mesothelioma (LATTICe-M), consisting of 512 patients and 3,446 resection WSIs ^23^. For benchmarking, we additionally included the publicly available Cancer Genome Atlas (TCGA)-mesothelioma cohort ^28^ used previously in the HPL study, which, although substantially smaller (75 patients, 84 images), provides another established external validation set.

### 5.1 Model Architecture and Pipeline

#### HPL mathematical formulation and validation setup

The HPL evaluation pipeline consists of three main stages: (i) tile-to-embedding extraction using the Barlow Twins–pretrained encoder, (ii) graph-based Leiden clustering followed by assignment of tiles to clusters, and (iii) construction of patient-level compositional representations with centred log-ratio (CLR) transformation for downstream predictive modelling.

##### Notation

Let *S* denote the set of WSIs in the biopsy cohort. For a WSI *s* ∈ *S* we extract a set of non-overlapping tiles:

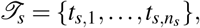

where each tile *t* is a 224 × 224 pixles patch at 5× (pixel size ≈ 1.8 *µ*m).

##### Encoder and tile embeddings

Let *f*_*θ*_ : *I* → ℝ^*D*^ be Barlow Twins encoder (Appendix A) with *D* = 128. For each tile *t*, its embedding is *z* = *f*_*θ*_ (*t*) ∈ ℝ^*D*^, optionally *ℓ*_2_-normalised. The encoder was trained on the LATTICe-M dataset, and the resulting pretrained weights were directly used to embed all tiles in the biopsy cohort for inference and evaluation.

##### HPC discovery and assignation

A set of histomorphological phenotype clusters (HPCs) *C* = *hpc*_1_, …, *hpc*_*c*_ was derived from the biopsy cohort by constructing a *k*-nearest-neighbour graph on a 250,000-tile subsample of embeddings and applying Leiden community detection. Given embeddings 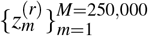, pairwise distances are computed and each node is connected to its *k* = 250 nearest neighbours, producing an adjacency matrix *W*. The Leiden algorithm then identifies a partition *P* that maximises modularity with a specific resolution *γ* (here we picked *γ* = 3.0). For the rest of the tiles in the cohort, its embedding *z* is assigned to the nearest cluster centroid *µ*_*j*_ using Euclidean distance:

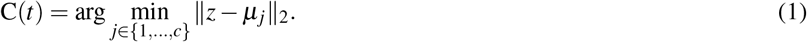

This produces, for each WSI *s*, cluster counts *n*_*s, j*_ = |{*t* ∈ *T*_*s*_ : C(*t*) = *j*}|. To ensure a strictly inductive setting, Leiden clustering and the fitting of the cluster centroids were performed exclusively on the feature vectors derived from the training set. The centroids were then fixed and applied to transform the validation features. All images underwent extensive quality control and preprocessing to ensure sufficient tissue content for downstream inference.

##### Subtype classification

For epithelioid vs. non-epithelioid classification, we use the frequency of HPCs per WSI to form a slide-level compositional vector. For each slide *s*, let *x*^(*s*)^ = clr(**a**^(*s*)^) ∈ ℝ^*c*^, where **a**^(*s*)^ is the HPC frequency vector and clr denotes the centred log-ratio transform (Appendix B). A logistic regression model is then fitted to predict the binary subtype label *y*^(*s*)^ ∈ {0, 1}.

The predicted probability of the epithelioid subtype is:

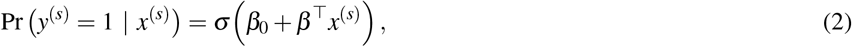

where *σ* (*u*) = (1 + *e*^−*u*^)^−1^. The parameters (*β*_0_, *β*) are estimated by minimising the regularised negative log-likelihood with an *ℓ*_1_ penalty weighted by *λ*_*ℓ*1_ = 3.5 after careful hyper-tuning of the model.

##### Survival modelling (Cox proportional hazards)

For patient-level survival analysis, we construct the CLR-transformed HPC composition vector *x*^(*i*)^ for each patient. Let *T* ^(*i*)^ denote the observed time and *δ* ^(*i*)^ the event indicator. A Cox proportional hazards model is then fitted:

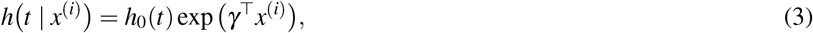

where *γ* are the log-hazard coefficients. Parameters are estimated by maximising the penalised partial likelihood:

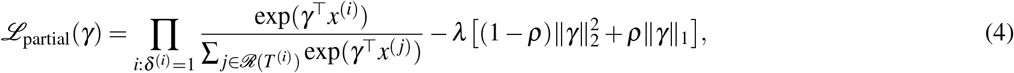

with *ℛ*(*t*) the risk set at time *t*, and a regularisation penalty controlled by *λ* = 0.5 and elastic net mixing parameter *ρ* = 0.3. The baseline hazard *h*_0_(*t*) is estimated non-parametrically using Breslow’s method ^29^, with ties handled via Efron’s method ^30^. Hazard ratios 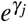 with 95% confidence intervals are reported. Patients are stratified into high and low-risk groups based on the cohort median risk, and Kaplan–Meier plots illustrate the corresponding survival probabilities.

##### Performance metrics and evaluation protocol

Validation used a patient-wise five-fold cross-validation ensuring strict separation of patients between folds (no tile or WSI from the same patient appears in both training and test splits). For classification we report Area under the curve (AUC), F1, sensitivity, and precision Scores, by showing the mean ± standard deviation across folds. For survival, we report Harrell’s concordance index (c-index) with bootstrap 95% confidence intervals. Where applicable, we test coefficient significance using Wald tests and perform permutation testing (shuffling labels) to evaluate the null distribution of metrics. We explored multiple clustering resolutions and identified resolution 3.0 as optimal, yielding the most distinct and stable clusters in terms of downstream performance. Regularisation parameters for both the logistic regression (*ℓ*_1_ penalty) and the penalised Cox proportional hazards model were systematically optimised through a grid search across candidate ranges, with the final configurations selected based on model stability and predictive performance.

##### Implementation details and reproducibility

Embeddings were computed using the publicly available HPL Encoder ^27^. We adopted the ResNet backbone as the feature encoder. This encoder was originally trained and validated on the LATTICe-M dataset and subsequently validated on the TCGA mesothelioma cohort in a previous study ^23^. The encoder was trained on a single Nvidia A100 GPU for approximately 48-72 hours over 60 epochs, using 800,000 training tile samples and 400,000 samples each for testing and validation. In the present work, we followed the same validation strategy, as in the TCGA-Meso cohort. We directly used the trained encoder weights to extract feature representations from biopsy tiles, without performing any additional fine-tuning or retraining (Figure 1g). Although, unlike the TCGA-Meso validation, which relied on a predefined dictionary of histomorphological clusters derived from resection specimens, we re-clustered the feature embeddings from scratch. This approach allowed us to identify biopsy-specific phenotypic clusters that more accurately reflect the morphological landscape of the biopsy domain. Final cluster count *c* is reported in the Results section. All experiments were run with fixed random seeds and patient stratification to ensure reproducibility.

## 6 Author Contributions

F.S. led the project, performed the analysis, and wrote the manuscript. F.D. and R.S. managed sample curation and metadata collection. F.F., S.L., and J.LQ performed histological annotation of the HPCs. A.S. provided expert annotation for the curated data and clinical coordination via the NETMESO network. S.L., J.LQ., and K.Y. supervised the study, provided critical guidance, and edited the manuscript. All authors reviewed and approved the final text.

## 7 Acknowledgments

We express our sincere gratitude to all pathologists and clinical contributors of the NETMESO network for their valuable expertise, case contributions, and ongoing support in annotation and case discussions. We also thank MESOBANK, French Mesothelioma Biobank, Centre Léon Bérard, Lyon, France, for its contribution to this research. We acknowledge the Cancer Research UK Scotland Institute for funding F.S. and for its continued support of the mesothelioma research program.

## 8 Supplementary Figures

**Figure S1:**
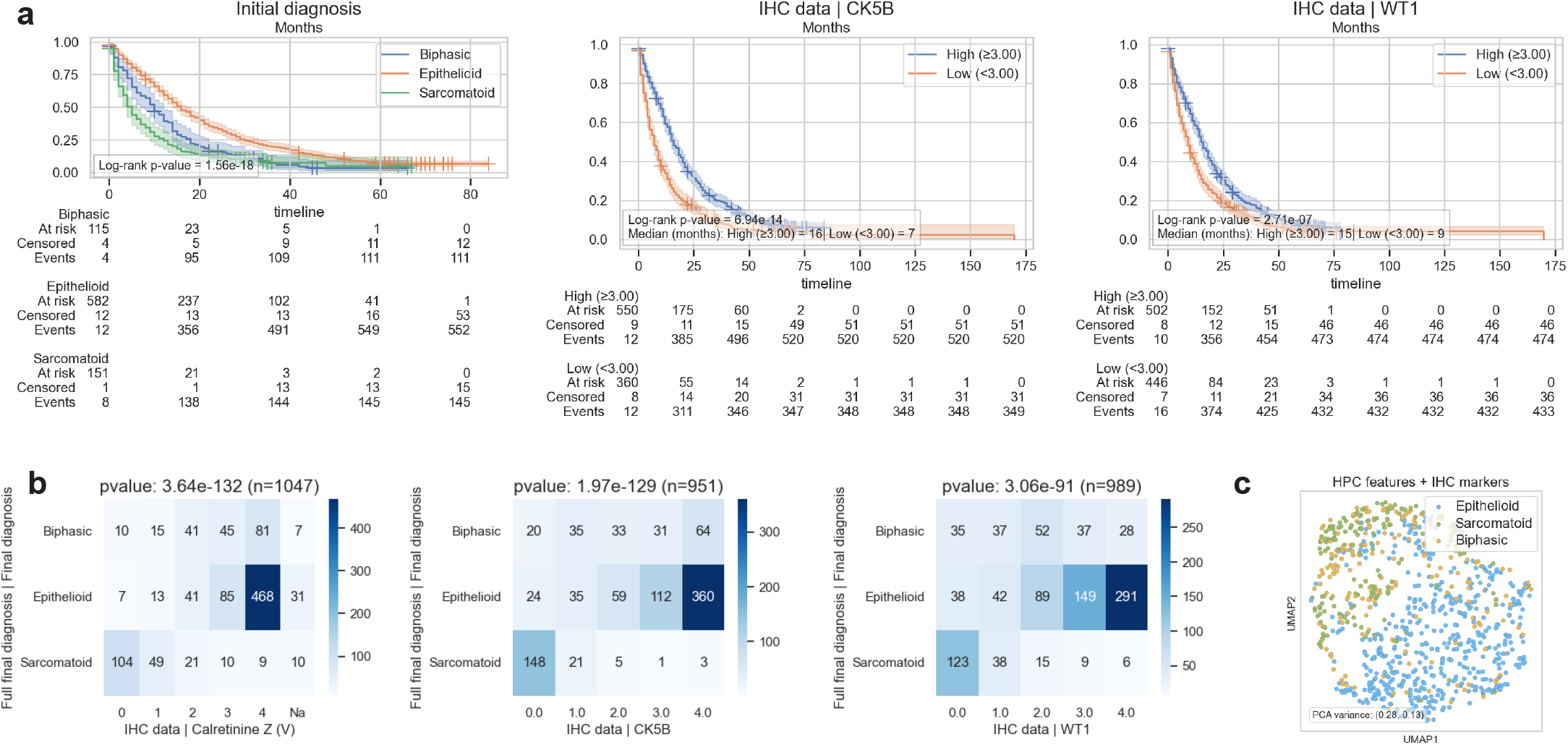
**(a)** Kaplan–Meier plot showing the significance of IHC markers and initial diagnosis in patient stratification. **(b)** Heatmaps showing the correlation between different IHC marker values and the three mesothelioma histopathological subtypes. **(c)** Patient sample UMAP coloured by their subtypes.

**Figure S2:**
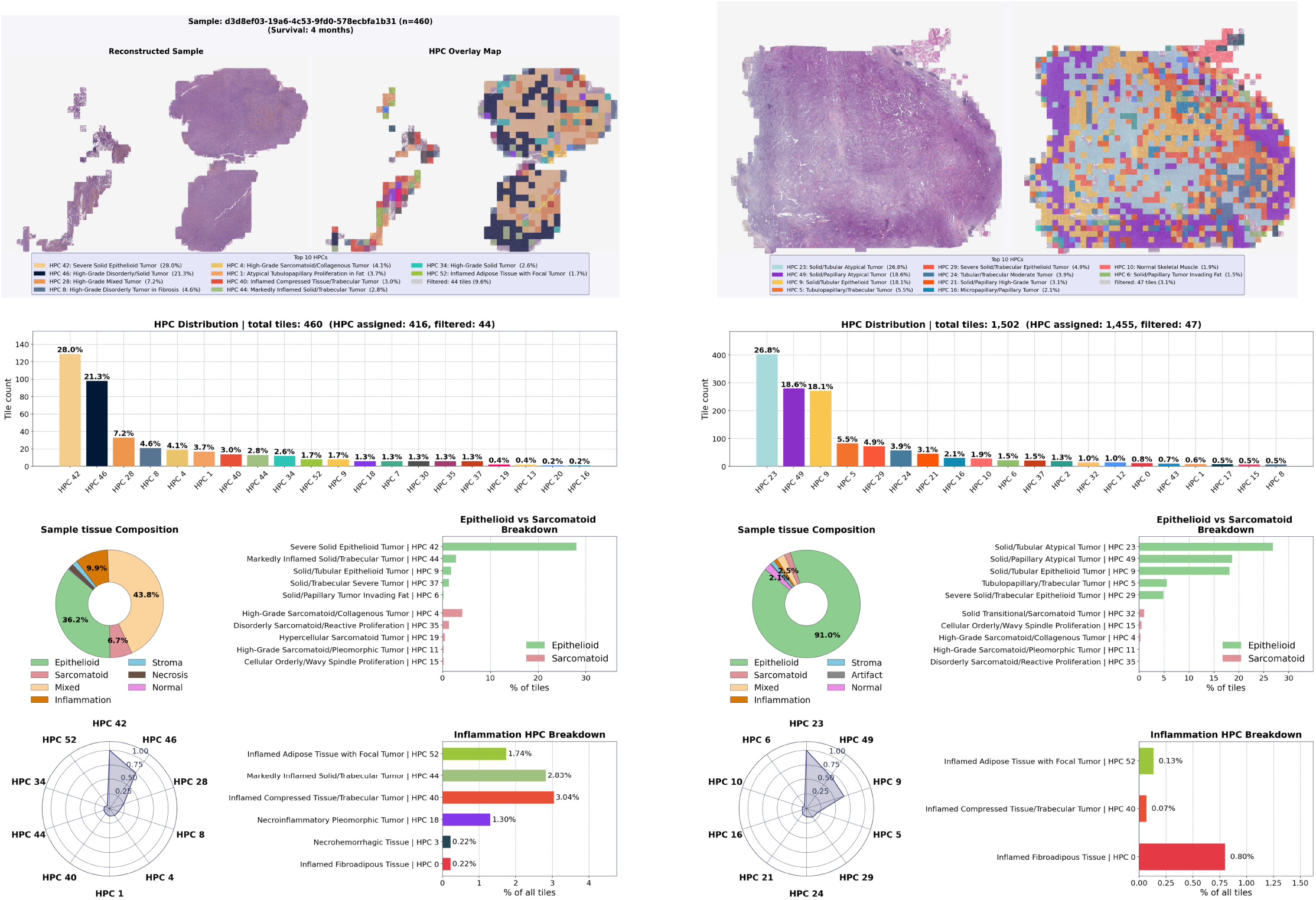
HPC breakdown of two samples from a high-survival patient and a low-survival patient, based on pathologist annotations for each HPC indicating tumor subtype, inflammation, and other histomorphological patterns.

**Figure S3:**
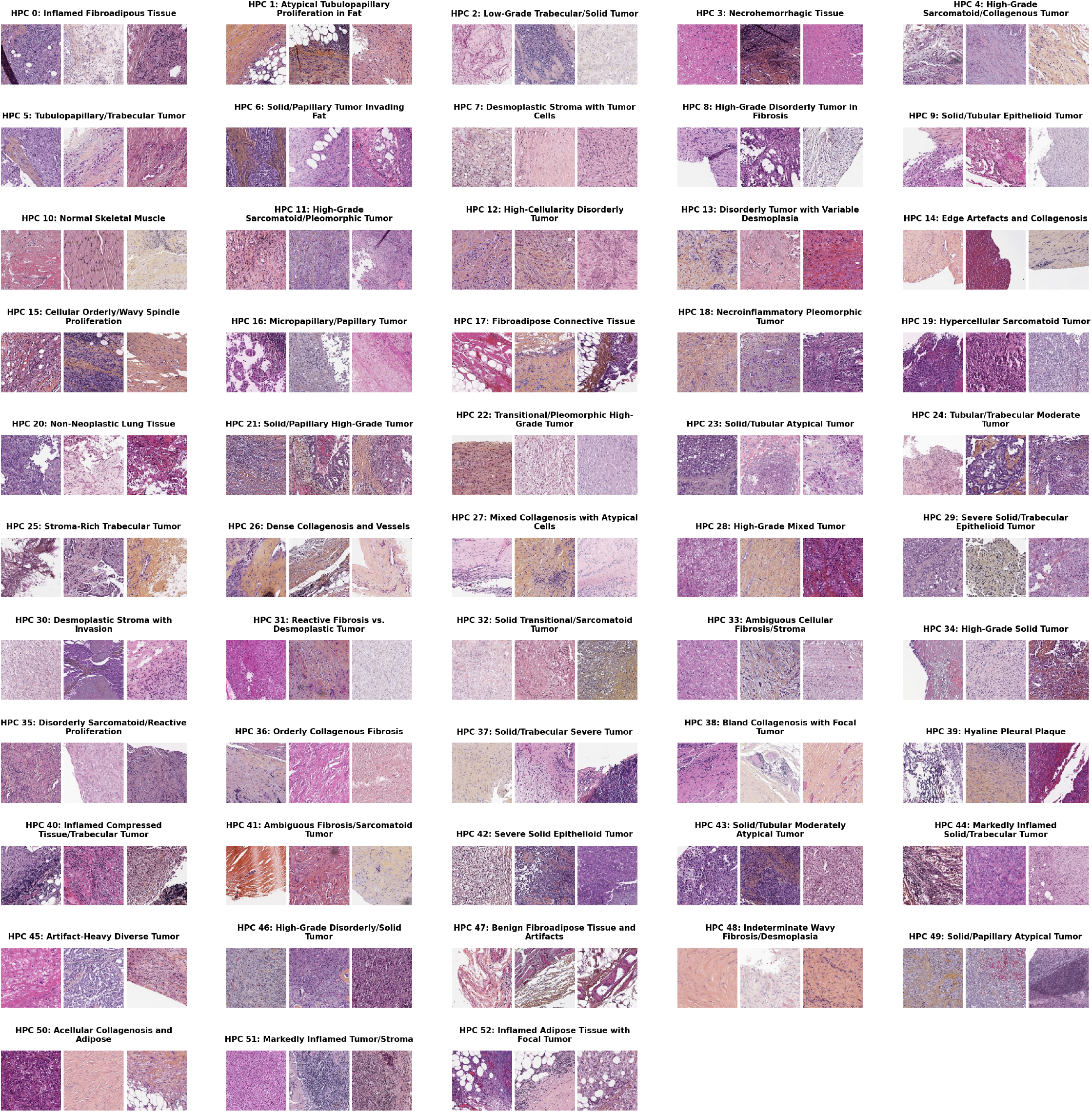
HPC Samples and their titles

## A Barlow Twins Training Formulation

During training, two stochastic augmentations (views) *v* and *v*^′^ are generated for each tile, producing embeddings *z* = *f*_*θ*_ (*v*) and *z*^′^ = *f*_*θ*_ (*v*^′^). For a mini-batch of size *N*, the empirical cross-correlation matrix *C* ∈ ℝ^*D*×*D*^ is defined as:

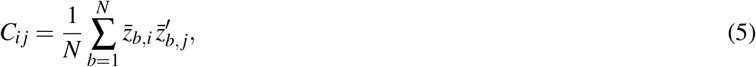

where 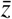 and 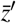 are the per-dimension mean-centred, variance-normalised embeddings across the batch. The Barlow Twins objective encourages invariance through diagonal alignment and decorrelation through off-diagonal suppression:

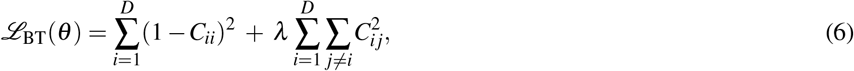

with *λ >* 0 controlling the off-diagonal penalty.

### Inference

After training, each tile *t* is embedded via *z* = *f*_*θ*_ (*t*) and used in the downstream HPL pipeline as described in the main text.

## B Compositional Vectors

We defined the raw HPC frequency vector for WSI *s* as

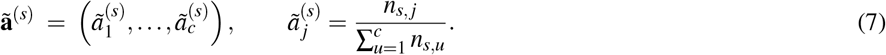

By construction **ã**^(*s*)^ is compositional: 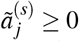 and 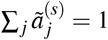.

These vectors can contain zeros; we apply multiplicative replacement prior to log-ratio transformations. We then transform each compositional vector **a**^(*s*)^ into Euclidean space via the centered log-ratio:

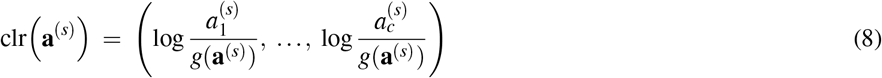

where 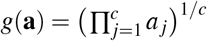 is the geometric mean. The clr vector has a zero sum: ∑ _*j*_ clr(**a**) _*j*_ = 0.

## Notes

### Competing Interest Statement

The authors have declared no competing interest.

### Summary of Updates

Small changes in the script including a few figure labels and grammer/spelling/wordings in the main texts

