## Supplementary Data and Figures for "Self-Supervised AI Discovery of Histomorphological Phenotypes from Routine Mesothelioma Biopsies"

Diagnosis and prognostic assessment in mesothelioma rely mainly on small tissue biopsies; however, most computational pathology models , including HPL-Meso [23], are trained on large tissue samples from surgical resections, thereby training data. However, these large resection specimens contain extensive morphologically diverse samples of tumour, stroma, necrosis, inflammation, and background tissues, whereas biopsies are much smaller, more uniform in composition, are prone to different processing artefacts (eg crush, over- vs under-fixation), sampling limitations. These differences create a significant mismatch between the data used for model development and the material used in routine diagnostic workflows, with direct implications for model reliability and clinical applicability.

| Variable | Value |
|---|---|
| Total cases | 1047 |
| Age | 22-100 (y) |
| Patient Survival | 0-170 (m) |

| Variable | Groups | Count (Percentage) |
|---|---|---|
| Sex | Female, Male | 241 (23.0%), 806 (77.0%) |
| Vital status | Alive, Dead | 91 (8.7%), 956 (91.3%) |
| Final diagnosis | Biphasic | 199 (19.0%) |
| | Epithelioid | 645 (61.6%) |
| | Sarcomatoid | 203 (19.4%) |
| FISH CDKN2A | Heterozygous Deletion | 10 (1.0%), |
| | Homozygous Deletion | 54 (5.2%), |
| | No Deletion | 22 (2.1%) |
| BAP1 | Lost, Retained | 545 (52.1%), 486 (46.4%) |
| Weight loss | No, Yes | 5 (0.5%), 87 (8.3%) |
| Pleural plaques | No, Yes | 142 (13.6%), 80 (7.6%) |
| History of cancer | No, Yes | 161 (15.4%), 205 (19.6%) |
| Type of therapy | Chemotherapy | 108 (10.3%) |
| | Radiotherapy | 53 (5.1%) |
| | Surgery | 21 (2.0%) |
| | Immunotherapy | 17 (1.6%) |

**Table 1:** Dataset Characteristics

### 3 Results

#### 3.1 Evaluation across a Multi-Center French Biopsy Cohort

For this study, we collected a large multicentre cohort of Hematoxylin-Eosin-Saffron (HES)- and Hematoxylin-Phloxine-Saffron (HPS)-stained biopsy slides received from pathology laboratories across several regions in France. All slides were reviewed by the NETMESO MESOPATH network pathologists, and digitised at the National Reference Centre in Lyon using Leica AT2 or Leica GT scanners at 20X magnification. The final dataset comprised 1,047 patients and 1,062 biopsy images. Among the patients, 77% ($n = 806$) were male and 23% ($n = 241$) were female. Patient ages ranged from 22 to 100 years, with a mean age of 76.5 years and a standard deviation of 8.7 years (Table 1). Treatment information was available for 199 patients, of whom 108 received chemotherapy, 53 radiotherapy, 21 surgery, and 17 immunotherapy. Both age and mesothelioma subtype were significantly ($p < 0.05$) associated with survival (Supplementary Figure S1). The cohort included 61.6% epithelioid ($n = 645$) and 38.4% non-epithelioid ($n = 402$) cases. Figure 1a illustrates the geographical distribution of referring institutes and highlights the top 10 contributing cities, led by Lyon. The map partly reflects differences in the number and capacity of pathology laboratories between regions.

In Figure 1b, we computed pairwise correlations between immunohistochemistry markers that are available in more than 100 samples. Significant correlations ($p < 0.01$) are indicated with an asterisk. CK5/6, calretinin, and WT1, which are established mesothelial lineage markers, showed strong positive correlations with one another. This mesothelial marker group also correlated positively with epithelial and cytokeratin markers, including CK7, AE1/AE3, and EMA. We also performed BerEP4, TTF1 and P40 and PAX8 as negative markers. A complete summary of reported marker distributions is provided in Supplementary Figure S2.

The median age at diagnosis for male patients was 79 years (minimum 44 years), compared with 77 years for female patients (minimum 22 years) (Figure 1c). Survival distributions differed substantially across mesothelioma subtypes, with mean survival times of 8.7, 13.2, and 21.6 months for sarcomatoid, biphasic, and epithelioid cases, respectively. Sarcomatoid cases exhibited the poorest prognosis, with a median survival of only 4 months, compared with 10 months for biphasic and 15 months for epithelioid cases (Figure 1d).

To highlight the diagnostic challenges associated with mesothelioma, Figure 1e shows that 36 patients initially diagnosed with epithelioid disease were later reclassified as biphasic following central review by the French national board of subspecialty experts. Among 72 cases with uncertain initial diagnoses, the final diagnoses changed to epithelioid ($n = 38$), sarcomatoid ($n = 17$), and biphasic ($n = 17$). In addition, three cases initially considered benign were ultimately classified as sarcomatoid malignancy, further emphasising the practical value of expert board review. Indeed, the Cohen's kappa score between initial and final diagnoses was 0.59, indicating only moderate agreement. Agreement scores between individual experts and the final consensus diagnosis ranged from 0.73 to 0.91 (Figure 1f).

#### 3.2 Discovery of Patient-Agnostic Histomorphological Phenotype Clusters (HPCs)

To characterise morphological patterns across the cohort, we used the resection specimen-trained feature encoder from [23] to generate tile-level feature embeddings and construct histomorphological phenotype clusters (HPCs) (Figure 1g). Leiden clustering identified 53 distinct HPCs. The UMAP projection of tile embeddings is shown in Figure 1h, with tile embeddings coloured according to cluster assignment. Although the encoder was trained exclusively on H&E images, it remained robust to staining variation and generated discriminative feature representations despite the presence of saffron staining. Representative clusters demonstrated clear morphological distinctions: cluster 20 exhibited pulmonary alveolar architectures; clusters 44 and 29 displayed progressively denser cellular patterns; cluster 52 was enriched for inflamed and fibrotic adipose tissue from the chest wall; and cluster 47 corresponded to stroma-dominated regions with collagen highlighted by the characteristic orange saffron stain.

As shown in Figure 1i-j, assigning biopsy tile embeddings to previously defined resection-derived HPCs (from the HPL-Meso study [23]) constrained the representation space to the morphological vocabulary of surgical resection samples. Consequently, several essentially resection-specific HPCs, such as HPC 28 (pleural plaque) and HPC 2 (compressed lung), were absent in biopsy specimens. In contrast, biopsy-derived HPCs exhibited a substantially more balanced distribution of tiles across clusters. Because this set of HPCs is discovered in the biopsy latent representations' domain and were therefore used for all downstream analyses effectively. The distribution of HPC contributions across cities is shown in Figure 1k and demonstrates a well-balanced and geographically diverse HPC set, and no HPC belongs to a single institute. Every HPC contained biopsy tiles originating from at least 42% of represented cities, while 50 of the 53 clusters received contributions from more than 50% of cities. Among the 107 represented cities, cluster 52 showed the lowest geographical coverage, with contributions from 45 cities, whereas cluster 12 showed the broadest coverage, with tiles originating from 99 cities. Figure 1l further illustrates the distribution of HPCs across individual patients. Of the 53 identified HPCs, 49 were represented in at least 20% of cases, while 23 HPCs were observed in more than 50% of cases, suggesting the presence of recurrent and common mesothelioma morphological patterns across the cohort.

Figure 2g shows the significant correlations between CLR-transformed HPC embeddings and immunohistochemical markers. Statistical significance was assessed using FDR-corrected p-values ($p < 0.05$), and each HPC was labelled according to the expert consensus annotations. The classical diagnostic markers of malignant mesothelioma, EMA CK5/6 and WT1, and to a less universal degree AE1/AE3, identify a set of clusters dominated by malignant epithelioid morphology, while sarcomatoid clusters are notably related to the loss of these markers, and frequently (though far from universally) to positivity for GATA3.

##### 3.4 HPCs Predict Core Mesothelioma Subtypes, BAP1 Loss, and Tumor Grade

Using the CLR-transformed HPC frequency vectors as case-level representations, we evaluated the ability of the learned histomorphological features to predict multiple downstream clinical and molecular variables, including histological subtype (epithelioid vs non-epithelioid), initial diagnosis, sarcomatoid versus biphasic subtype, BAP1 loss status, nuclear grading, and immunohistochemistry marker expression such as CK5/6 and Calretinin. Performance metrics, including Area under the curve (AUC), F1-score, sensitivity, and precision, are summarised in Table 2.

The framework achieved a cross-validated average AUC of 0.94 for distinguishing epithelioid from non-epithelioid cases ($n = 975$), demonstrating that the unsupervised HPC representations captured highly discriminative subtype-associated morphology. HPCs 16 (papillary architecture), 24 (tubular architecture), and 25 (trabecular architecture), all characterised by moderate nuclear atypia and minimal inflammation, were among the most significant predictors of epithelioid disease. In contrast, HPC 28, characterised

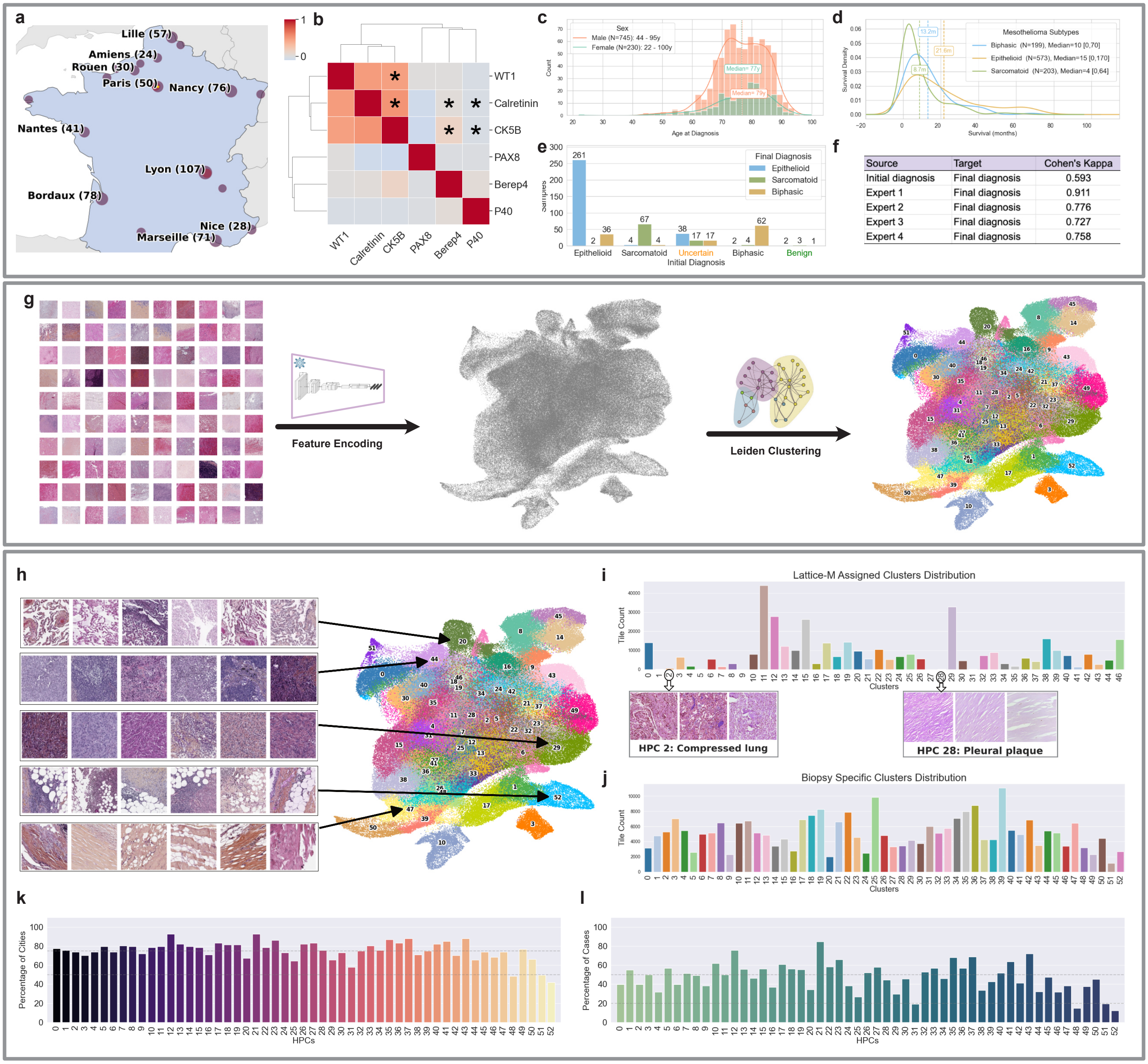


**Figure 1:** Overview of the multicentre mesothelioma biopsy cohort, immunohistochemical marker relationships, diagnostic variability, and histomorphological phenotype clusters (HPCs). **(a)** Geographical distribution of referring pathology centres and patient case numbers across French cities. **(b)** Pairwise correlation matrix of immunohistochemistry (IHC) markers, with significant correlations indicated by asterisks (p<0.01). **(c–d)** Distribution of patient age at diagnosis and survival across mesothelioma subtypes. **(e–f)** Diagnostic variability analysis showing changes between initial and final diagnoses and agreement scores between pathologists and consensus diagnosis. **(g–h)** Workflow for generating biopsy-derived HPCs and UMAP visualisation of tile embeddings coloured by cluster assignment. Representative clusters illustrate distinct morphological patterns, including alveolar tissue, inflamed and highly cellular tumorous tissue, and adipose tissue. **(i–j)** Comparison between resection-derived and biopsy-specific HPC assignments, highlighting the limited representation of certain resection-associated morphologies in biopsy samples. **(k–l)** Geographical and patient-level distribution of biopsy-derived HPCs, demonstrating broad representation of clusters across centres and cases.

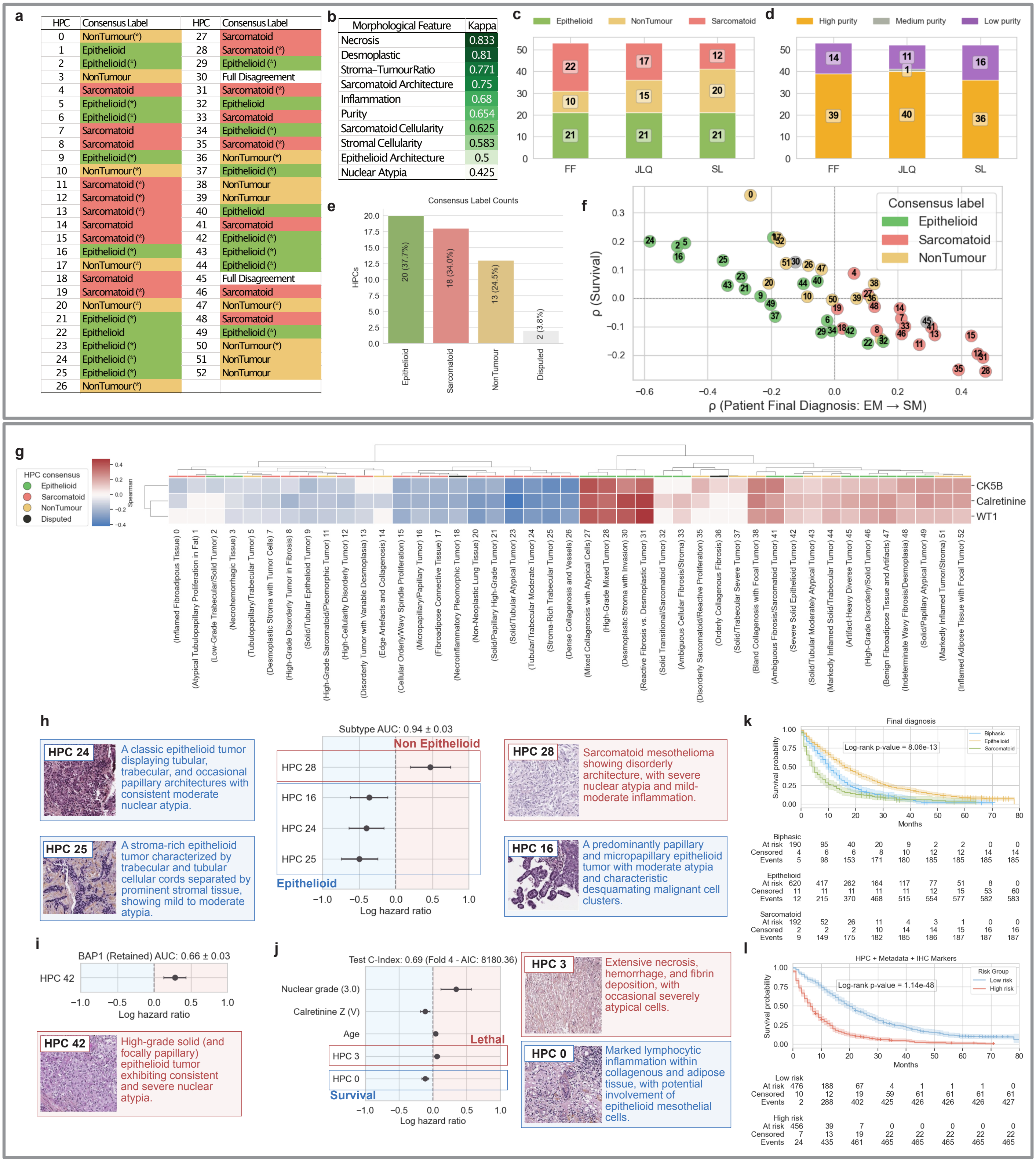


**Figure 2:** HPC Annotations and downstream tasks (a-f) HPC annotations' analysis (g)Significant IHC marker correlations with HPCs. (h-j) Significant HPCs associated with downstream tasks. (K,I) Kaplan-Meier survival curves stratified by histological subtype and by integrated risk groups derived from clinical, molecular, and HPC features.

| Task | Classes (n) | ROC-AUC | F1 Score | Sensitivity | Precision |
|---|---|---|---|---|---|
| Subtype | Epitheliod, Non-epithelioid (975) | 0.94 ± 0.03 | 0.87 ± 0.04 | 0.86 ± 0.04 | 0.87 ± 0.04 |
| Initial Diagnosis | Biphasic, Epithelioid, Sarcomatoid, Uncertain (975) | 0.91 ± 0.02 | 0.77 ± 0.02 | 0.76 ± 0.02 | 0.78 ± 0.02 |
| Non-Epithelioid | Sarcomatoid, Biphasics (402) | 0.79 ± 0.04 | 0.72 ± 0.04 | 0.72 ± 0.04 | 0.73 ± 0.04 |
| BAP-1 | Retained, Lost (959) | 0.66 ± 0.03 | 0.62 ± 0.03 | 0.62 ± 0.03 | 0.62 ± 0.03 |
| Calretinin | 0,1,2,3,4 (928) | 0.73 ± 0.02 | 0.45 ± 0.03 | 0.41 ± 0.03 | 0.53 ± 0.04 |
| CK5/6 | 0, 1, 2, 4 (884) | 0.70 ± 0.02 | 0.42 ± 0.03 | 0.40 ± 0.03 | 0.47 ± 0.04 |
| Nuclear grade | 1,2,3 (643) | 0.59 ± 0.04 | 0.42 ± 0.04 | 0.42 ± 0.04 | 0.44 ± 0.04 |

**Table 2:** Performance of downstream classification tasks, including histological subtype, BAP1 loss, nuclear grading, and marker expression prediction using CLR-transformed HPC embeddings.

by disorderly sarcomatoid architecture and severe nuclear atypia, was the strongest predictor of non-epithelioid cases (Figure 2h). These findings indicate that the HPC representation space captures biologically meaningful architectural transitions associated with mesothelioma subtype progression.

The model also demonstrated predictive capability for BAP1 loss, achieving an average AUC of 0.66 with a sensitivity of 0.62 ($n = 959$). HPC 42, characterised by epithelioid solid architecture with severe nuclear atypia and mild-to-moderate inflammation, was significantly associated with BAP1 loss in case-level predictions. This observation is consistent with previous studies reporting higher frequencies of BAP1 loss in epithelioid mesothelioma compared with other histological subtypes [24], [25]. Overall, these results suggest that the self-supervised HPC representations encode not only morphological subtype information, but also clinically relevant molecular and phenotypic characteristics directly from routine biopsy images.

##### 3.5 High Grade Tumour Histological Phenotypes Drive Patient Risk

We next evaluated the prognostic value of the learned HPC information using survival analysis. Clinical variables, including age, sex, and histological subtype diagnosis (epithelioid, sarcomatoid, and biphasic), achieved an average test-set concordance index (C-index) of 0.65. Additionally, immunohistochemistry markers, including Calretinin, CK5/6, and WT1 (positive markers), demonstrated a predictive value with a 5 fold cross validation average C-index of 0.61 for the test set.

Incorporating histomorphological information derived from HPC statistics (such as Pielou's evenness index and Shannon diversity) alongside their frequency of occurrence in the patient samples increased the test-set C-index to 0.65 (Table 3). Also, filtering out certain HPCs using pathologist annotations (such as restricting the analysis to tumourous HPCs ($n = 38$) or high-purity HPCs ($n = 40$)) did not improve the C-index. However, combining the HPC-derived frequencies and statistical metrics with both IHC data from positive markers and clinical information significantly increased performance to a cross-validated average C-index of 0.68.

| Variables | Train Concordance Index | Test Concordance Index |
|---|---|---|
| Clinical (Age, Sex, Mesothelioma subtype diagnosis) | 0.65 ± 0.00 | 0.65 ± 0.02 |
| Immunohistochemistry markers | 0.61 ± 0.01 | 0.61 ± 0.03 |
| HPCs (HPC statistics) | 0.68 ± 0.01 | 0.65 ± 0.02 |
| Tumorous (Epithelioid and Sarcomatoid) HPCs | 0.65 ± 0.01 | 0.63 ± 0.02 |
| High Purity HPCs (%75) | 0.67 ± 0.01 | 0.64 ± 0.02 |
| HPCs + Immunohistochemistry + Clinical Data | 0.71 ± 0.00 | 0.68 ± 0.02 |

**Table 3:** Survival prediction performance using clinical variables, immunohistochemistry markers, and HPC-derived features individually and in combination. Reported metrics are averaged across 5-fold cross-validation and are presented as mean ± standard deviation.

In summary, our results show that models trained on resection specimens can still achieve clinically meaningful accuracy when applied to biopsy material. Notably, the biopsy validation also strengthens the evidence for the robustness of the HPL pipeline, even with a smaller cohort than LATTICe-M. Also, despite being learned without diagnostic supervision, the HPC features showed associations with molecular characteristics and immunohistochemical marker expression. Importantly, HPC-derived features provided prognostic information comparable to established clinical and molecular variables and further improved survival prediction when integrated with them. The identified HPCs corresponded to interpretable architectural and cytological patterns recognised by expert pathologists, suggesting that the learned representation space reflects genuine disease morphology rather than dataset-specific visual signals.

**Notation.** Let $\mathscr{S}$ denote the set of WSIs in the biopsy cohort. For a WSI $s \in \mathscr{S}$ we extract a set of non-overlapping tiles:

$$\mathscr{T}_s = \{t_{s,1}, \ldots, t_{s,n_s}\},$$

where each tile $t$ is a $224 \times 224$ pixles patch at $5\times$ (pixel size $\approx 1.8\ \mu$m).

**Encoder and tile embeddings.** Let $f_\theta : \mathscr{I} \to \mathbb{R}^D$ be Barlow Twins encoder (Appendix A) with $D = 128$. For each tile $t$, its embedding is $z = f_\theta(t) \in \mathbb{R}^D$, optionally $\ell_2$-normalised. The encoder was trained on the LATTICe-M dataset, and the resulting pretrained weights were directly used to embed all tiles in the biopsy cohort for inference and evaluation.

**HPC discovery and assignation** A set of histomorphological phenotype clusters (HPCs) $\mathscr{C} = \{hpc_1, \ldots, hpc_c\}$ was derived from the biopsy cohort by constructing a $k$-nearest-neighbour graph on a 250,000-tile subsample of embeddings and applying Leiden community detection. Given embeddings $\{z_m^{(r)}\}_{m=1}^{M=250{,}000}$, pairwise distances are computed and each node is connected to its $k = 250$ nearest neighbours, producing an adjacency matrix $W$. The Leiden algorithm then identifies a partition $\mathscr{P}$ that maximises modularity with a specific resolution $\gamma$ (here we picked $\gamma = 3.0$). For the rest of the tiles in the cohort, its embedding $z$ is assigned to the nearest cluster centroid $\mu_j$ using Euclidean distance:

$$\mathrm{C}(t) = \arg \min_{j \in \{1,\ldots,c\}} \|z - \mu_j\|_2. \tag{1}$$

This produces, for each WSI $s$, cluster counts $n_{s,j} = |\{t \in \mathscr{T}_s : \mathrm{C}(t) = j\}|$. To ensure a strictly inductive setting, Leiden clustering and the fitting of the cluster centroids were performed exclusively on the feature vectors derived from the training set. The centroids were then fixed and applied to transform the validation features. All images underwent extensive quality control and preprocessing to ensure sufficient tissue content for downstream inference.

**Subtype classification.** For epithelioid vs. non-epithelioid classification, we use the frequency of HPCs per WSI to form a slide-level compositional vector. For each slide $s$, let $x^{(s)} = \mathrm{clr}(\mathbf{a}^{(s)}) \in \mathbb{R}^c$, where $\mathbf{a}^{(s)}$ is the HPC frequency vector and clr denotes the centred log-ratio transform (Appendix B). A logistic regression model is then fitted to predict the binary subtype label $y^{(s)} \in \{0, 1\}$.

The predicted probability of the epithelioid subtype is:

$$\Pr\left(y^{(s)} = 1 \mid x^{(s)}\right) = \sigma\left(\beta_0 + \beta^\top x^{(s)}\right), \tag{2}$$

where $\sigma(u) = (1 + e^{-u})^{-1}$. The parameters $(\beta_0, \beta)$ are estimated by minimising the regularised negative log-likelihood with an $\ell_1$ penalty weighted by $\lambda_{\ell_1} = 3.5$ after careful hyper-tuning of the model.

**Survival modelling (Cox proportional hazards).** For patient-level survival analysis, we construct the CLR-transformed HPC composition vector $x^{(i)}$ for each patient. Let $T^{(i)}$ denote the observed time and $\delta^{(i)}$ the event indicator. A Cox proportional hazards model is then fitted:

$$h\big(t \mid x^{(i)}\big) = h_0(t)\exp\big(\gamma^\top x^{(i)}\big), \tag{3}$$

where $\gamma$ are the log-hazard coefficients. Parameters are estimated by maximising the penalised partial likelihood:

$$\mathscr{L}_{\text{partial}}(\gamma) = \prod_{i:\delta^{(i)}=1} \frac{\exp(\gamma^\top x^{(i)})}{\sum_{j\in\mathscr{R}(T^{(i)})}\exp(\gamma^\top x^{(j)})} - \lambda\left[(1-\rho)\|\gamma\|_2^2 + \rho\|\gamma\|_1\right], \tag{4}$$

with $\mathscr{R}(t)$ the risk set at time $t$, and a regularisation penalty controlled by $\lambda = 0.5$ and elastic net mixing parameter $\rho = 0.3$. The baseline hazard $h_0(t)$ is estimated non-parametrically using Breslow's method [29], with ties handled via Efron's method [30]. Hazard ratios $e^{\gamma_j}$ with 95% confidence intervals are reported. Patients are stratified into high and low-risk groups based on the cohort median risk, and Kaplan–Meier plots illustrate the corresponding survival probabilities.

### 8 Supplementary Figures

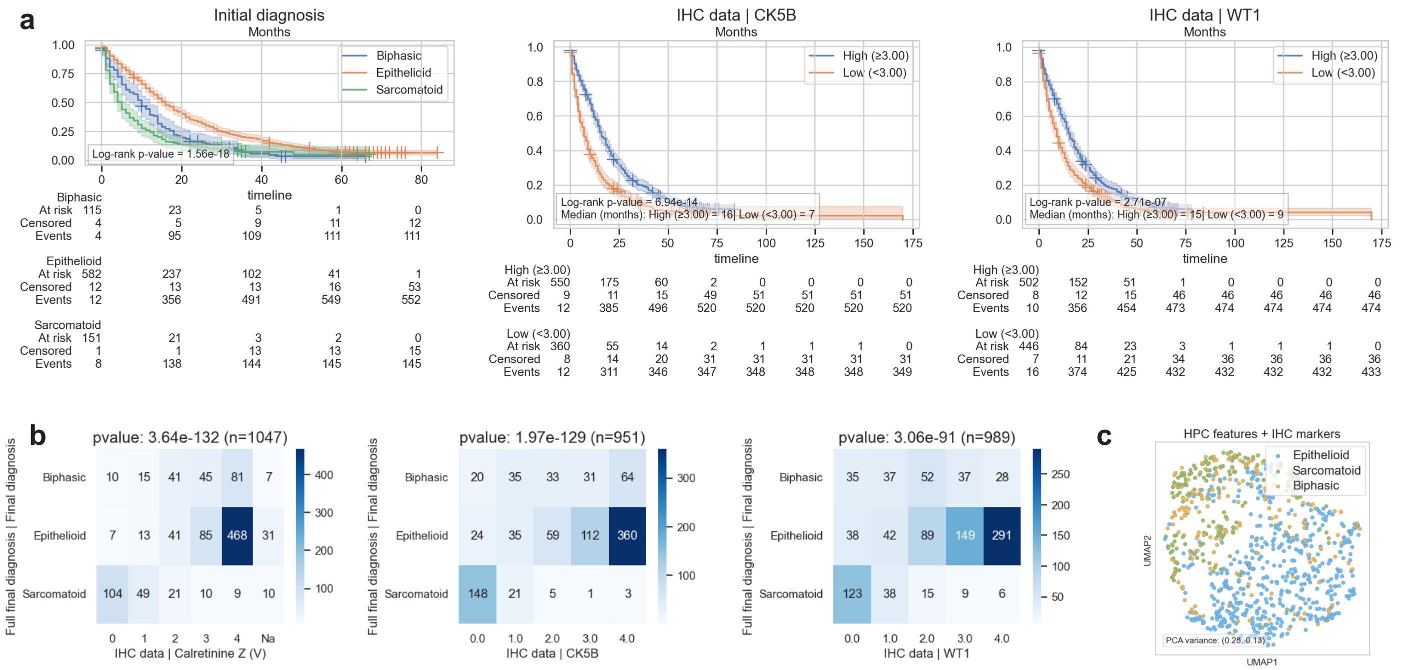


**Figure S1:** (**a**) Kaplan–Meier plot showing the significance of IHC markers and initial diagnosis in patient stratification. (**b**) Heatmaps showing the correlation between different IHC marker values and the three mesothelioma histopathological subtypes. (**c**) Patient sample UMAP coloured by their subtypes.

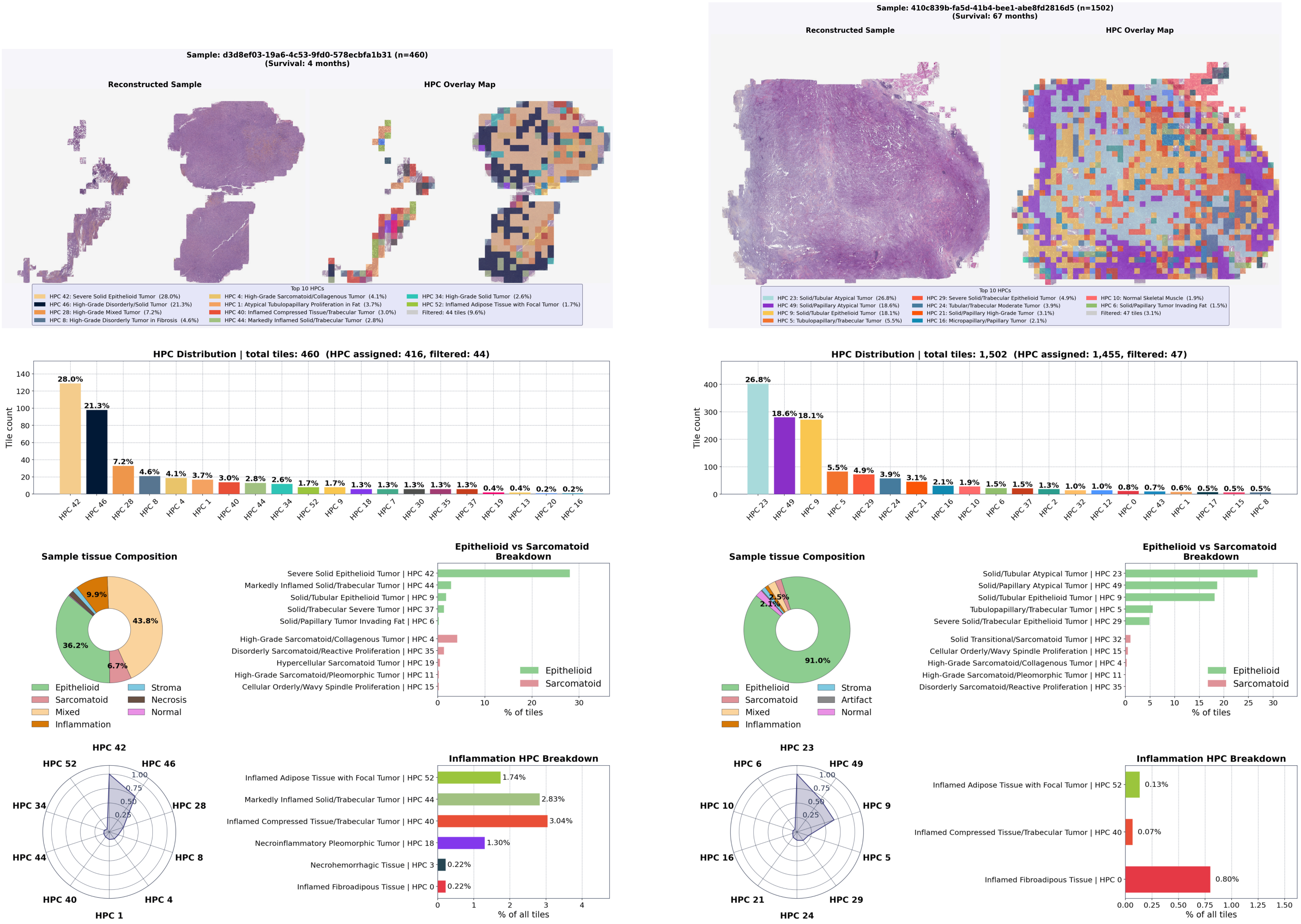


**Figure S2:** HPC breakdown of two samples from a high-survival patient and a low-survival patient, based on pathologist annotations for each HPC indicating tumor subtype, inflammation, and other histomorphological patterns.

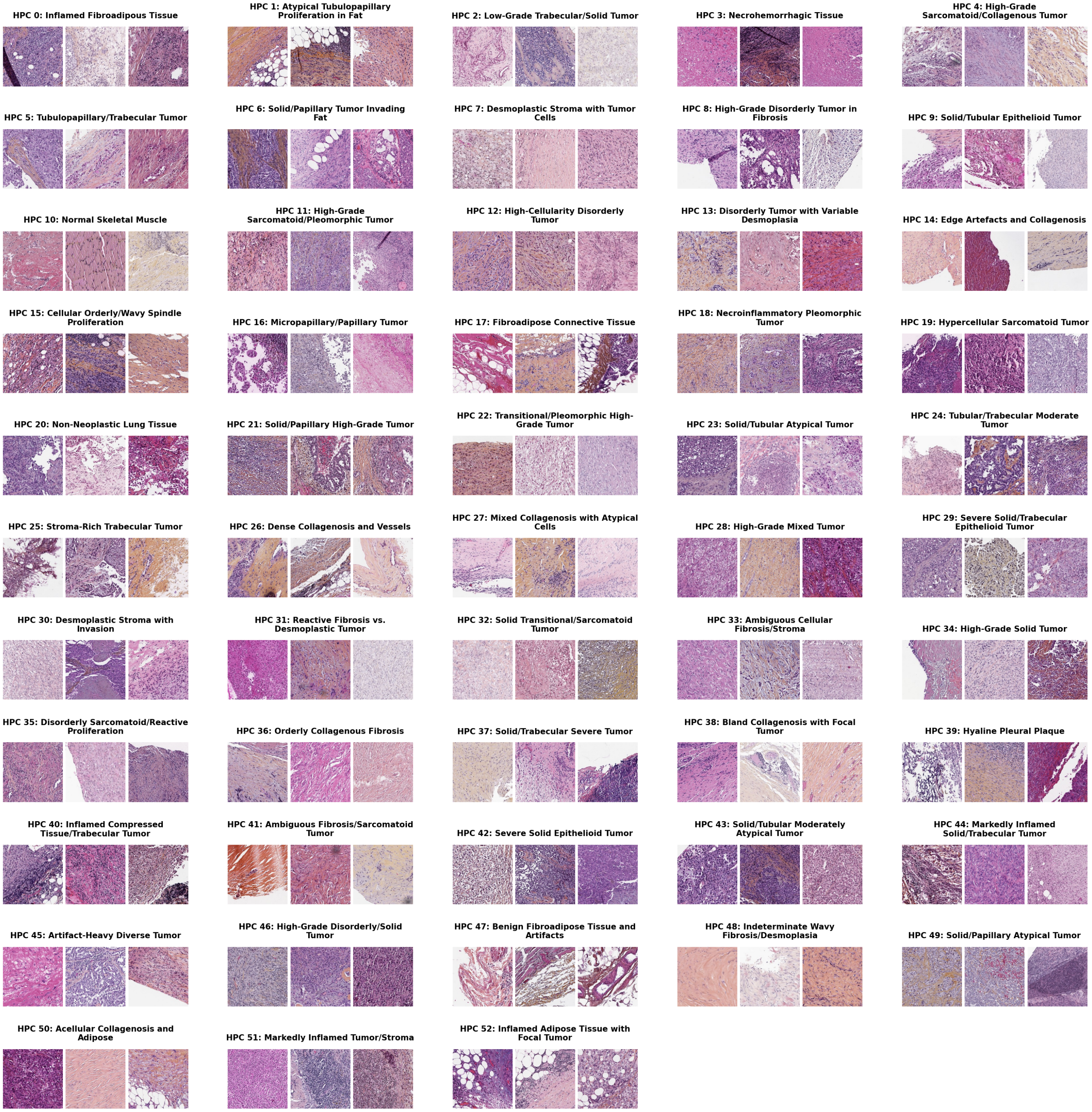


**Figure S3:** HPC Samples and their titles

### A Barlow Twins Training Formulation

During training, two stochastic augmentations (views) $v$ and $v'$ are generated for each tile, producing embeddings $z = f_\theta(v)$ and $z' = f_\theta(v')$. For a mini-batch of size $N$, the empirical cross-correlation matrix $C \in \mathbb{R}^{D \times D}$ is defined as:

$$C_{ij} = \frac{1}{N} \sum_{b=1}^{N} \bar{z}_{b,i} \, \bar{z}'_{b,j}, \tag{5}$$

where $\bar{z}$ and $\bar{z}'$ are the per-dimension mean-centred, variance-normalised embeddings across the batch. The Barlow Twins objective encourages invariance through diagonal alignment and decorrelation through off-diagonal suppression:

$$\mathscr{L}_{\mathrm{BT}}(\theta) = \sum_{i=1}^{D} (1 - C_{ii})^2 \; + \; \lambda \sum_{i=1}^{D} \sum_{j \neq i} C_{ij}^2, \tag{6}$$

with $\lambda > 0$ controlling the off-diagonal penalty.

**Inference.** After training, each tile $t$ is embedded via $z = f_\theta(t)$ and used in the downstream HPL pipeline as described in the main text.

### B Compositional Vectors

We defined the raw HPC frequency vector for WSI $s$ as

$$\tilde{\mathbf{a}}^{(s)} \; = \; \left(\tilde{a}_1^{(s)}, \ldots, \tilde{a}_c^{(s)}\right), \qquad \tilde{a}_j^{(s)} = \frac{n_{s,j}}{\sum_{u=1}^{c} n_{s,u}}. \tag{7}$$

By construction $\tilde{\mathbf{a}}^{(s)}$ is compositional: $\tilde{a}_j^{(s)} \geq 0$ and $\sum_j \tilde{a}_j^{(s)} = 1$.

These vectors can contain zeros; we apply multiplicative replacement prior to log-ratio transformations. We then transform each compositional vector $\mathbf{a}^{(s)}$ into Euclidean space via the centered log-ratio:

$$\operatorname{clr}\left(\mathbf{a}^{(s)}\right) \; = \; \left( \log \frac{a_1^{(s)}}{g(\mathbf{a}^{(s)})}, \; \ldots, \; \log \frac{a_c^{(s)}}{g(\mathbf{a}^{(s)})} \right) \tag{8}$$

where $g(\mathbf{a}) = \left(\prod_{j=1}^{c} a_j\right)^{1/c}$ is the geometric mean. The clr vector has a zero sum: $\sum_j \operatorname{clr}(\mathbf{a})_j = 0$.
